# Metagenomic Data Reveal Type I Polyketide Synthase Distributions Across Biomes

**DOI:** 10.1101/2023.01.09.523365

**Authors:** Hans W. Singh, Kaitlin E. Creamer, Alexander B. Chase, Leesa J. Klau, Sheila Podell, Paul R. Jensen

## Abstract

Microbial polyketide synthase (PKS) genes encode the biosynthesis of many biomedically important natural products, yet only a small fraction of nature’s polyketide biosynthetic potential has been realized. Much of this potential originates from type I PKSs (T1PKSs), which can be delineated into different classes and subclasses based on domain organization and structural features of the compounds encoded. Notably, phylogenetic relationships among PKS ketosynthase (KS) domains provide a method to classify the larger and more complex genes in which they occur. Increased access to large metagenomic datasets from diverse habitats provides opportunities to assess T1PKS biosynthetic diversity and distributions through the analysis of KS domain sequences. Here, we used the webtool NaPDoS2 to detect and classify over 35,000 type I KS domains from 137 metagenomic data sets reported from eight diverse biomes. We found biome-specific separation with soils enriched in modular *cis*-AT and hybrid *cis*-AT KSs relative to other biomes and marine sediments enriched in KSs associated with PUFA and enediyne biosynthesis. By extracting full-length KS domains, we linked the phylum Actinobacteria to soil-specific enediyne and *cis*-AT clades and identified enediyne and monomodular KSs in phyla from which the associated compound classes have not been reported. These sequences were phylogenetically distinct from those associated with experimentally characterized PKSs suggesting novel structures or enzyme functions remain to be discovered. Lastly, we employed our metagenome-extracted KS domains to evaluate commonly used type I KS PCR primers and identified modifications that could increase the KS sequence diversity recovered from amplicon libraries.

**Importance:** Polyketides are a crucial source of medicines, agrichemicals, and other commercial products. Advances in our understanding of polyketide biosynthesis coupled with the accumulation of metagenomic sequence data provide new opportunities to assess polyketide biosynthetic potential across biomes. Here, we used the webtool NaPDoS2 to assess type I PKS diversity and distributions by detecting and classifying KS domains across 137 metagenomes. We show that biomes are differentially enriched in KS domain classes, providing a roadmap for future biodiscovery strategies. Further, KS phylogenies reveal both biome-specific clades that do not include biochemically characterized PKSs, highlighting the biosynthetic potential of poorly explored environments. The large metagenome-derived KS dataset allowed us to identify regions of commonly used type I KS PCR primers that could be modified to capture a larger extent of KS diversity. These results facilitate both the search for novel polyketides and our understanding of the biogeographical distribution of PKSs across earth’s major biomes.

## Introduction

Microorganisms are a valuable source of structurally diverse specialized metabolites including many with clinically relevant biological activities (1–2). Recent advances in DNA sequencing technologies and molecular genetics have fostered new discovery paradigms based on the mining of natural product biosynthetic gene clusters (BGCs) from microbial genomes (3). Instrumental to this field are online tools such as antiSMASH 5.0 (4) and PRISM (5) that detect and classify BGCs within query data. Additionally, repositories like MIBiG (6), which list BGCs that have been experimentally linked to compounds, and IMG-ABC (7), which details BGCs within sequenced microbial genomes, serve as important comparison points for genome mining efforts.

Polyketides represent a major class of pharmaceutically relevant microbial specialized metabolites (8) and an important target for genome mining research. Their biosynthesis is mediated by polyketide synthase (PKS) genes, which can be classified into types I-III depending on their enzymatic domain structure (8). Type I PKSs are composed of multi-domain proteins and represent the largest source of polyketide natural products within the MIBiG repository (6). A minimal type I PKS requires an acetyltransferase (AT) domain, which selects the appropriate building block, an acyl carrier protein (ACP) domain, to which the building block is tethered, and a ketosynthase (KS) domain, which catalyzes chain elongation between the growing polyketide and the ACP-bound extender unit (9–11). Based on the organization and function of these domains, type I PKS genes can be delineated into three classes (12), the first of which functions as a multi-modular assembly line (referred to here as modular *cis*-AT) where each KS domain catalyzes one round of chain elongation. The second class generally has only one module (monomodular) with the KS domain functioning iteratively to catalyze more than one round of chain elongation (12), while the third class has a modular organization that uses stand-alone AT domains (*trans*-AT) that occur outside of the PKS gene (13).

It has long been recognized that KS phylogenies can be used to distinguish sequences associated with type I modular *cis*-AT, iterative, and *trans*-AT PKSs and thus make broader predictions about the types of PKS genes in which they occur (14–17). Type I KS phylogenies can further provide insight into the types of compounds produced (e.g., enediynes, PUFAs, NRPS-PKS hybrids) or the functional role of the KS in polyketide assembly (e.g., loading vs. extension modules). The web tool NaPDoS2 automates this process by detecting and classifying KS domain sequences from genomic, metagenomic, or amplicon query data (14–15) and using these sequences to make broader predictions about PKS diversity. In this way, NaPDoS2 circumvents the need for complete PKS gene or BGC assembly, which can be particularly challenging for multimodular type I PKSs. This makes it ideal for assessing biosynthetic potential within poorly assembled metagenomic datasets.

While metagenomic data has provided important new insights into natural product biosynthetic diversity, the use of PCR to amplify KS domains from environmental DNA (eDNA) allows for the rapid detection of low frequency sequences. This approach has enabled the large-scale comparison of KS diversity across environmental samples (18–30) and guided the discovery of novel natural products from cloned eDNA (25). The KS primers used to assess T1PKS diversity were originally designed based on modular *cis*-AT KSs detected in the phyla Actinobacteria, Cyanobacteria, and Deltaproteobacteria (20–21). While this primer set has been slightly modified over the years (22–25), it is unclear how well it conforms to the KS diversity observed in metagenomic datasets. Recent evidence that all 18 KS domains extracted from understudied taxa would not be amplified by this primer set suggests modifications may be warranted (31).

Assessing biosynthetic diversity using metagenomic datasets carries distinct advantages over amplicons in that complete BGCs can be captured and PCR biases avoided. In addition, metagenome assembled genomes (MAGs) have revealed unknown or poorly studied microbial taxa from soils (31) and seawater (32) that are enriched in uncharacterized BGCs and thus new targets for natural product discovery. However, the fragmented nature of most metagenomic assemblies provides poor representation of rare community members, which are an important source of natural products (31). Indeed, a recent analysis of 1500 metagenomes binned on average only 5.3 MAGs per metagenome (33). To date, metagenomes have largely been used to analyze the biosynthetic potential of individual biomes with the aim of finding new products (31–32,34–36). For example, direct cloning of metagenomic DNA from the gut microbiome led to the discovery of new polyketides (34) while metagenomes from root endophyte microbiomes led to the identification of PKS products that play key roles in disease suppression (35). Metagenomes have also been used to link compounds to producers as demonstrated recently for a sponge metagenome (36). Comparisons of PKS diversity across biomes are less common, although a recent study suggested that specific chemistry is not limited or amplified by environment (37).

In this study, we used NaPDoS2 to detect and classify over 35,000 type I KS domains from 240 Gbps of metagenomic data derived from eight environmental biomes. Using KS phylogenies, we detected biome-specific clades that are distinct from those associated with experimentally characterized BGCs. Additionally, we show that less than 3% on average of the KS domains in each metagenome are associated with MAGs, supporting the value of these sequences as a proxy to assess biosynthetic diversity. Finally, access to diverse KS sequences provided an opportunity to evaluate the effectiveness of a widely used type I KS primer set.

## Results

### Type I PKS distributions across biomes

To survey the distribution of type I PKSs across eight environmental biomes, we used NaPDoS2 to identify KS domains in 137 shotgun metagenomes from the following sample types: forest/agricultural soil, rhizosphere, peat soil, freshwater, seawater, freshwater sediment, marine sediment, and host-associated (Table S1). In total, 35,116 KS domains associated with type I PKSs and 409 KS domains associated with type I FASs (fatty acid synthases) were detected across all datasets (minimum alignment length >200 aa). The NaPDoS2 output further delineated the non-FAS KS domains into three classes (*cis*-AT, iterative *cis*-AT, and *trans*-AT) and, for some, into eight subclasses (hybrid *cis*-AT, cis-loading module, olefin synthase, PUFA, enediyne, aromatic, PTM, hybrid *trans*-AT) (Table S2). To validate the NaPDoS2 KS classifications, we analyzed representative sequences across the range of KS domain subclasses by running the associated metagenomic contigs through antiSMASH (4). In each case, the KSs were associated with PKS genes that matched the NaPDoS2 classification (Fig. S1).

The majority of metagenome-extracted type I KSs (37.5%) were classified by NaPDoS2 as *cis*-AT with no further subclass designation. The next most abundant designations were KS identified as iterative *cis*-AT PUFA (20.9%) and modular *cis*-AT hybrid (18.9%) (Table S2). We also analyzed the MAGs binned from each metagenome through the JGI IMG pipeline finding that, on average, only 2.7% of the type I KS domains within a given metagenome were located within MAGs (Fig. S2). This highlights the utility of targeting KSs and the limitations of focusing on MAGs when assessing biosynthetic potential using metagenomic datasets.

A PCoA analysis based on a Bray-Curtis dissimilarity matrix showed a significant separation of biomes based on type I KS composition (PERMANOVA, p < 0.001, R_2_ = 0.499) with PUFA, *cis*-AT, and hybrid *cis*-AT KS domains representing major drivers of biome separation between marine and non-marine samples (Fig. 1a). To further address differences in KS composition across biomes, we determined the frequency of KSs per Gbp and found that marine sediments had significantly more PUFA and enediyne sequences (Fig. 1b) than other biomes (Tukey’s HSD, p< 0.01). Likewise, forest/agricultural soil and rhizosphere metagenomes encoded significantly more hybrid *cis*-AT KS domains per Gbp (p< 0.01), and forest/agricultural soil metagenomes encoded more *cis*-AT KS domains (p<0.01) than non-soil biomes (Fig. 1b).

**Figure 1.**
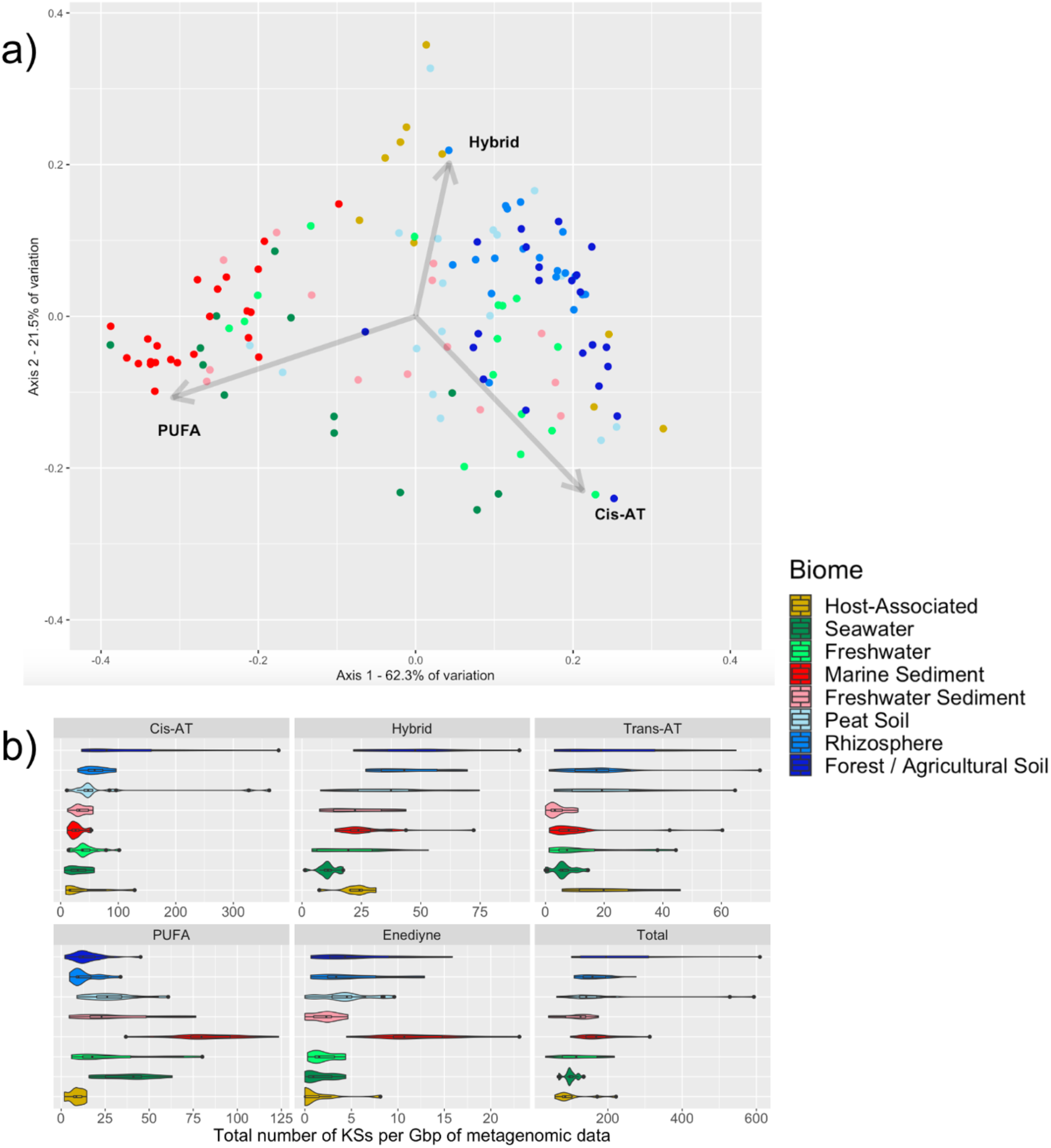
Biome-specific type I KS diversity and abundance. (A) PCoA of type I KS domain distributions after transformation using a Bray-Curtis dissimilarity matrix. Each point represents a metagenomic dataset (colored by biome). Arrows indicate the three KS domain types driving the most variation. (B) Violin plots showing the number of type I KS domains per Gbp of metagenomic data across eight biomes.

### KS richness and diversity across biomes

To compare KS richness and diversity across biomes using a more standardized framework, we focused on full-length KS domains that could be extracted from the metagenomes. Of the 7,945 full-length sequences recovered, 49.7% originated from soils and 50.3% from non-soil biomes. After clustering the full-length KS domains into operational biosynthetic units (OBUs) over a range of 70-95% (38), we found that soil and freshwater sediment biomes consistently carried greater KS richness based on Chao1 index values than marine sediment and seawater biomes at all clustering levels (Fig. S3). However, the only significant differences in richness were observed in forest/agricultural and peat soils, which were more diverse than non-soil biomes at the 90% and 95% clustering thresholds (p<0.01).

We next identified the number of KS domains in OBUs shared between biomes by rarefying each biome to 580 full-length KS sequences and performing pairwise comparisons (Fig. 2). Overall, marine sediment and seawater biomes shared the greatest number of OBUs at all clustering levels except 70%. Shared OBUs were also commonly identified in pairwise comparisons among forest/agricultural soil, peat soil, rhizosphere, and freshwater sediment biomes, and these always ranked among the top 10 in terms of the number of KS domains within the shared OBUs. Surprisingly, at 95% clustering, no OBUs were shared between freshwater and freshwater sediments and only the seawater and marine sediment biomes had more than 10 KS sequences within shared OBUs. In fact, at both the 95% and 90% clustering levels, many biome combinations (64% and 46%, respectively) contained no shared OBUs, with the maximum number of KS sequences within OBUs that were shared at 42 and 96, respectively. In contrast, at the lower clustering thresholds of 80% and 70%, all biome combinations shared at least two KS sequences, and the maximum number of shared KS sequences was 195 and 416, respectively (Fig. 2).

**Figure 2.**
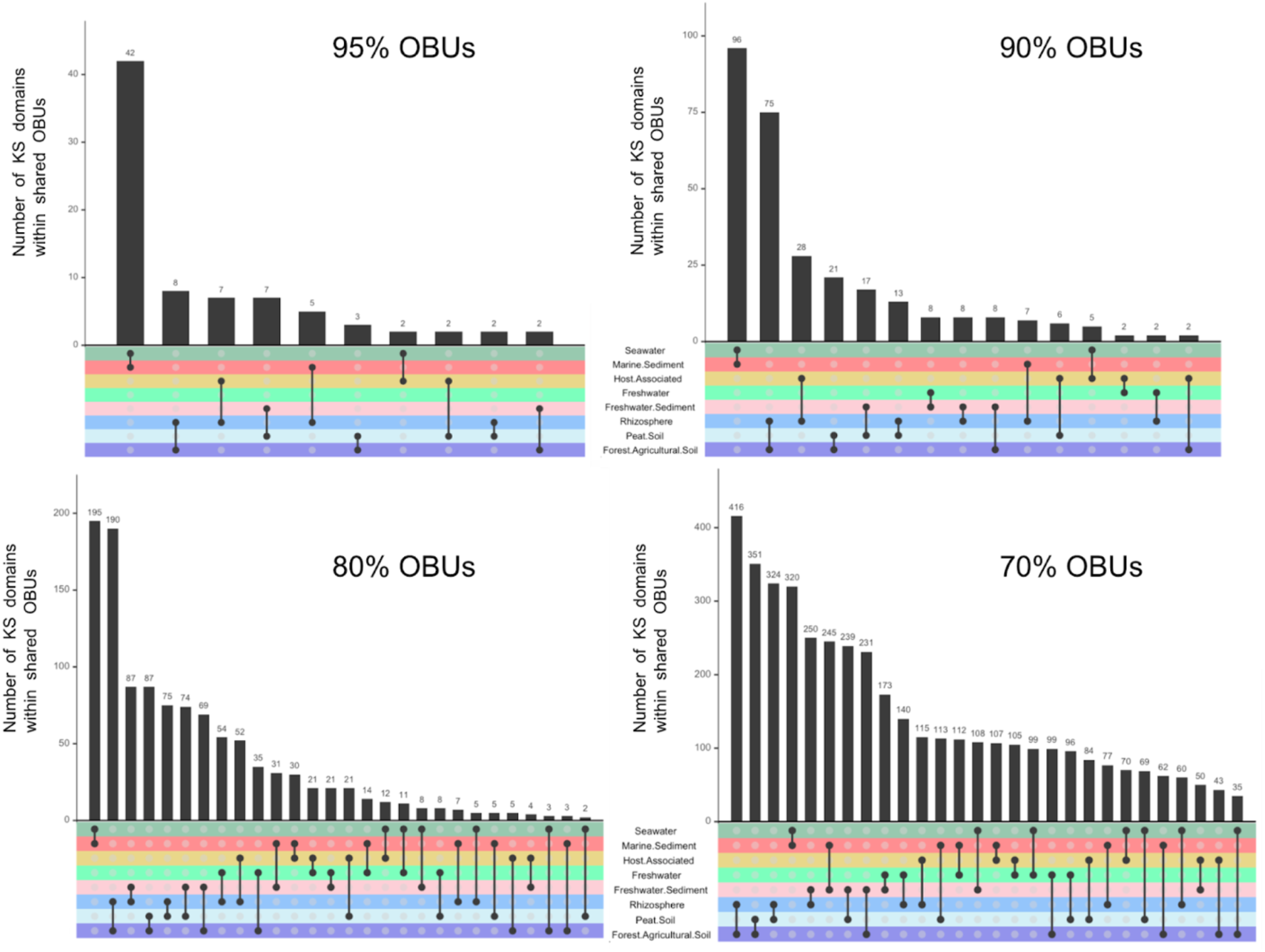
Shared KS diversity between biomes. The number of KS domain sequences (y-axis) in OBUs shared between two biomes (x-axis, black dots connected with a line) based on pairwise comparisons performed using OBU clustering values from 70-95% amino acid identity.

### Type I KS domains form five major groups

A sequence similarity network (SSN) was used to visualize relatedness among the 7,945 full-length metagenome-extracted KS domains in the context of their NaPDoS2 classification (Fig. S4). Additionally, 3,040 full-length KS domains identified in the MIBiG database (version 2.0) were included (2,149 *cis*-AT/iterative, 725 *trans*-AT, 126 hybrid *cis*-AT, 29 PUFA, and 11 enediyne KS domains). The hybrid *cis*-AT (n=1,746), *trans*-AT (n=831), PUFA (n=1,996), and enediyne (n=210) KS domains are clearly distinguished within the SSN. The *cis*-AT/iterative cluster (n=3,162) represents the fifth group and includes KSs classified by NaPDoS2 as modular (assembly line) *cis*-AT (including olefin synthase and loading module subclasses) and iteratively acting *cis*-AT (including iterative aromatic and iterative PTM subclasses). The three PUFA KS clusters correlate with the three KS domains that are usually found in PUFA PKSs (12,22). We next generated KS phylogenies to ask if biome-specific clades could be detected within each of the five major groups identified in the SSN.

### Biome-specific and uncharacterized clades within the *cis*-AT/iterative group

Since *cis*-AT KS domains were responsible for driving a large portion of the separation among biomes (Fig. 1), a phylogeny was constructed using the 3,162 metagenomic and 2,149 MIBiG sequences that comprised the *cis*-AT/iterative cluster in the SSN. This phylogeny revealed a large clade in which 98.0% of the 923 metagenome-extracted KS domains mapped to soil biomes (Fig. 3a, yellow inner ring). This clade includes more than 50% of the MIBiG reference sequences, all of which originate from multimodular, assembly-line *cis*-AT PKSs. Outside of this soil-dominant clade, the remaining 2239 metagenome-extracted KS domains within the cis-AT/iterative group were more evenly spread between soil (50.5%) and non-soil (49.5%) biomes. Additionally, the phylogeny revealed a small clade of 89 sequences that originated from sponge metagenomes (Fig. 3a, brown inner ring) and was exclusive of any MIBiG sequences. This is consistent with previous work describing the sponge-specific *sup* KS clade (29–30).

**Figure 3.**
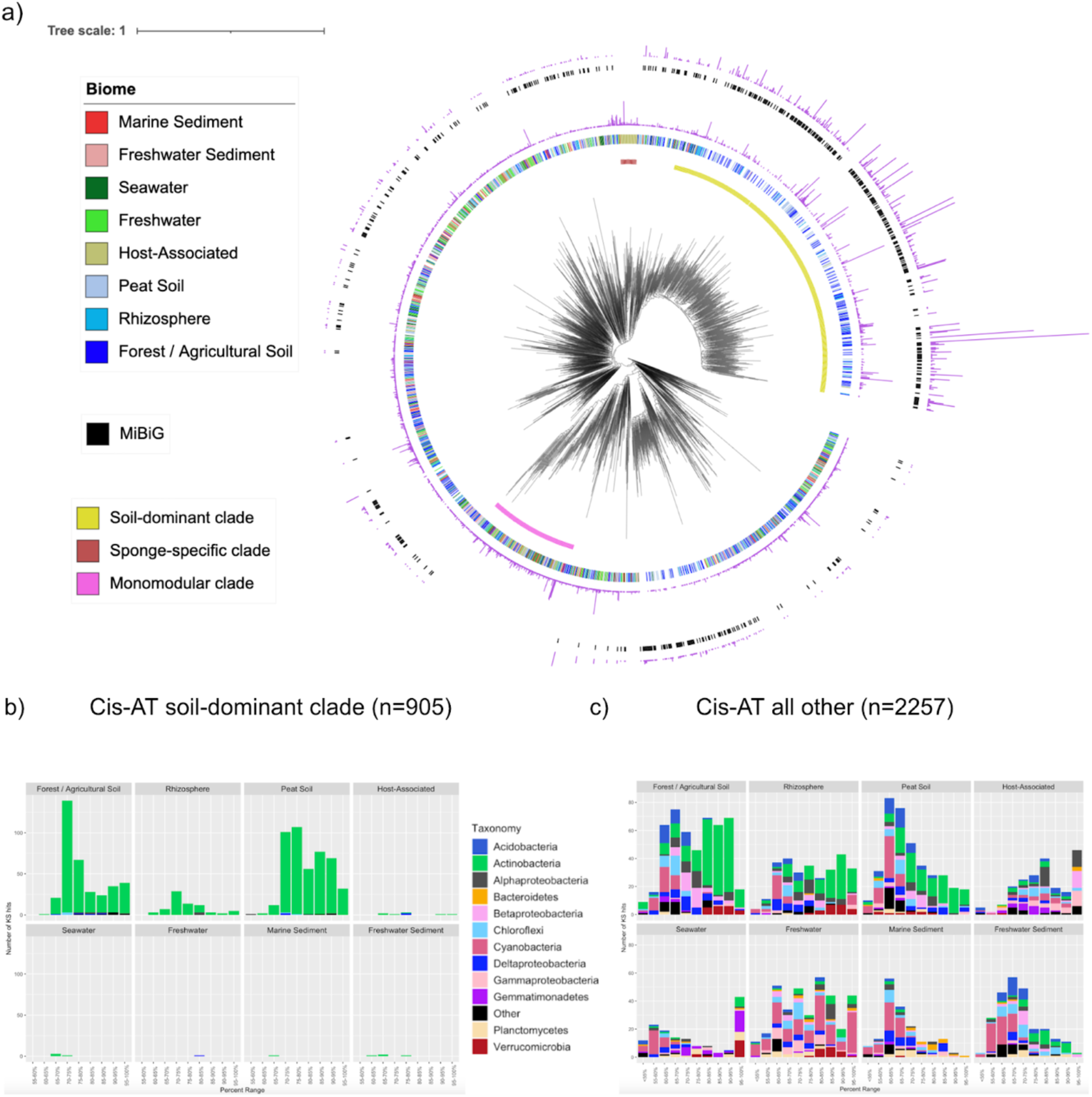
Phylogeny and taxonomic distribution of KS domains from the cis-AT/iterative group across biomes. (A) FastME phylogeny of full-length, metagenome-extracted, *cis-* AT/iterative group KS OBUs (70% sequence identity). The innermost ring includes a soil-dominant clade (n=905, yellow), a sponge-specific clade (n=89, brown), and a monomodular clade that does not include any MIBiG sequences (n=379, pink). The second ring indicates the biome from which each KS was derived and the third ring (purple) indicates the number of metagenome-extracted KS domains in each OBU. The fourth ring (black) depicts the MIBiG database *cis*-AT/iterative group KS domains grouped into OBUs, and the fifth ring (purple) shows the number of MIBiG-extracted KS sequences in each OBU. (B) Taxonomic distributions across eight biomes for the soil-dominant clade. (C) Taxonomic distributions across eight biomes for all *cis*-AT/iterative group KSs other than the soil dominant clade.

The taxonomic affiliations of the metagenome-extracted KS domains, assessed using the closest NCBI Blastp match, revealed that 96.2% of the *cis*-AT/iterative sequences within the soil-dominant clade could be assigned to the phylum Actinobacteria (Fig. 3b). The sequences outside this clade had a wider taxonomic distribution (Fig. 3c), with 23.5% assigned to Actinobacteria, 18.2% to Cyanobacteria, and 3-9% to nine low abundance phyla. In contrast with the soil-dominant clade, the sponge-specific clade displayed greater taxonomic diversity, with most sequences mapping to the phyla Gemmatimonadetes (24.7%) and Alphaproteobacteria (20.2%).

The KS phylogeny of the *cis*-AT/iterative group also revealed a large clade (379 metagenome-extracted KS domains) that did not group with any MIBiG sequences (Fig. 3a, pink inner ring), indicating a lack of functional characterization. The closest MIBiG KS domains were classified as iterative PTMs, which have been reported from Actinobacteria and Proteobacteria (6) and have wide-ranging antifungal activities (39) making them of interest for natural product discovery efforts. The PKS genes responsible for PTM biosynthesis contain a single module (monomodular) with the unique PKS-NRPS hybrid domain architecture of KS-AT-DH-KR-C-A-TE (39). To determine if the KSs in this clade were associated with PTM-like gene architectures, we searched the metagenomic assemblies and found two contigs of sufficient length for AntiSMASH analysis (Fig. S5). For the other KSs in this clade, we used BLAST to identify the top matching KSs (>70% sequence identity) in NCBI RefSeq and determined they were all associated with monomodular PKSs that possessed PTM-like architectures (Fig. S5). The RefSeq monomodular BGCs were observed in four phyla (Proteobacteria, Verrucomicrobia, Planctomycetes and Bacteroidetes) and encoded slightly different tailoring enzymes compared to the PTMs characterized to date in Actinobacteria and Proteobacteria. A multi-locus phylogeny showed that the metagenome extracted BGCs were closely related to RefSeq BGCs observed in Verrucomicrobia and Proteobacteria and distinct from the MIBiG PTM BGCs (Fig. S5). Further, the RefSeq and metagenome extracted BGCs from this clade had no MIBiG matches when analyzed for “known cluster hits”, indicating that considerable monomodular PKS diversity remains to be functionally characterized.

### Enediyne KS diversity across biomes

Given that marine sediments had significantly more enediyne KS sequences than other biomes (Fig. 1b), we assessed their novelty in comparison with enediyne KSs from the MIBiG database (n=10) and the NCBI RefSeq genome database (n=151). Enediynes represent a rare class of natural products that contain two acetylenic groups conjugated to a double bond within either a nine or 10-membered ring. They are highly cytotoxic, have been developed into effective cancer drugs (40), and to date have only been reported from the phylum Actinobacteria and marine ascidian extracts (41). A phylogeny generated using the 210 full-length, metagenome-extracted enediyne KS sequences revealed a soil-specific lineage (n=50) that co-localized with the genome-derived (NCBI RefSeq) sequences from Actinobacteria. This lineage encompassed all 10 MIBiG enediyne KS domains (Fig. 4). The remaining 160 metagenome-extracted enediyne KS domains were linked to a range of phyla including Cyanobacteria, Proteobacteria, Firmicutes, Bacteroidetes, Chloroflexi, and Spirochaetes based on their associations with the NCBI RefSeq sequences, with only two sequences of Actinobacterial origin. To further assess the non-Actinobacterial enediyne KS domains detected in RefSeq, we analyzed the respective genomes using AntiSMASH and found that all of the KS domains were associated with enediyne-like T1PKSs. Notably, 40% of the metagenome-extracted enediyne KS domains in the non-Actinobacterial portion of the phylogeny originated from marine sediments, with many (n=31) observed in a large, sediment-specific clade. Sequences in this clade shared 85% or greater amino acid similarity (NCBI BlastP) with a KS domain observed in a Deltaproteobacteria MAG (Fig. S6), suggesting a potentially new source of enediyne natural products. Interestingly, four of the RefSeq enediyne KS domains were observed in anaerobes (three from Deltaproteobacteria, and one from Spirochaetes), which are also not known to produce enediyne compounds.

**Figure 4.**
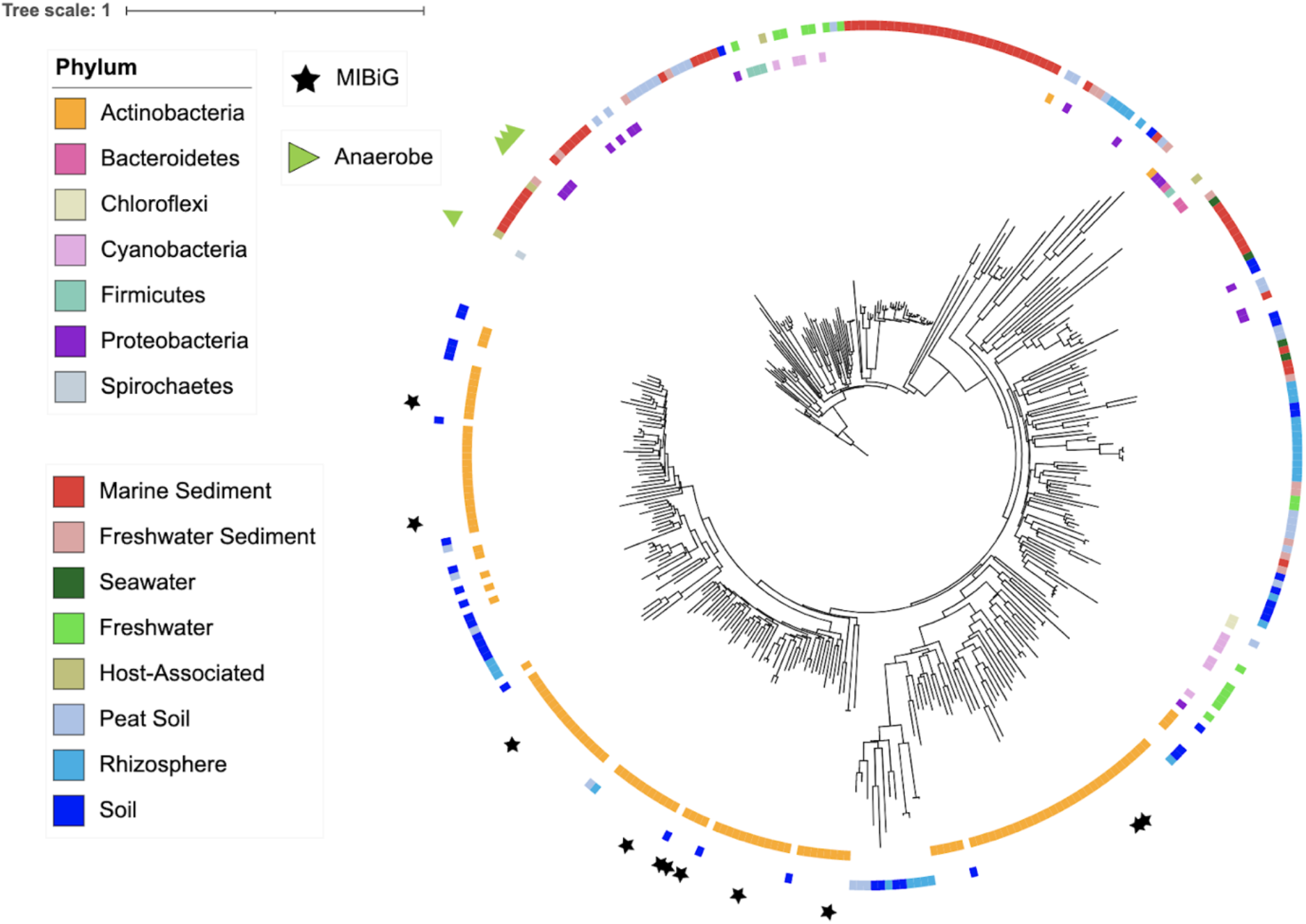
Distribution of enediyne KS domains across biomes and taxa. FastME phylogeny built using full-length enediyne KS domains obtained from metagenomes (n=210, outer ring, colored by biome), the NCBI RefSeq database (inner ring, colored by taxonomy), and the MIBiG database (stars).

### Hybrid *cis*-AT, *trans*-AT, and PUFA KS domains largely lack biome-specificity

We next investigated the taxonomic distributions and biome specificities of the metagenome-extracted hybrid *cis*-AT, *trans*-AT, and PUFA KS clusters identified in the SSN (Fig. S4). Hybrid *cis*-AT KS domains catalyze the condensation of an acyl group onto a PCP-tethered intermediate. *Trans*-AT KS domains occur in PKSs in which the AT domain acts in *trans* as a standalone acetyltransferase (13). PUFA BGCs occur across a wide range of bacterial phyla and are characterized by two to three KS domains (within modules *pfaA, pfaB* and *pfaC*) that collectively encode the production of linear carbon chains with multiple *cis* double bonds (22). We found no noticeable biome-specific clustering across the hybrid *cis*-AT (n=1746), *trans*-AT (n=831), and PUFA KS domains (n=1996) (Figs. S7-S9). Based on their top NCBI Blastp matches, PUFA KS domains were most often assigned to Deltaproteobacteria (27.7%), hybrid *cis*-AT KS domains to Cyanobacteria (24.7%), and *trans*-AT KS domains to Gammaproteobacteria (27.0%) and Firmicutes (21.7%). To date, very few PUFA pathways have been experimentally linked to a BGC, with previous work highlighting five known *pfaA* KS clusters (22) based on the production of eicosapentaenoic acid (EPA), docosahexaenoic acid (DHA), docosahexaenoic/docosapentaenoic acids (DHA-pseudo), arachidonic acid (AA), and heterocyst glycolipids (HGL). From our metagenome-extracted *pfaA* PUFA KS domains (n=1,170), over 92% fell outside these known clusters (Fig. S10), suggesting significant potential for the discovery and characterization of new PUFAs.

### Metagenomic KS diversity allows for the evaluation of KS primer sets

The metagenome-extracted KS domains analyzed here are diverse in terms of their taxonomic affiliations and biome of origin. Importantly, they are not biased towards cultured strains. As such, we saw an opportunity to evaluate existing type I KS primer sets using our metagenome-extracted sequences. The most commonly used type I KS primer set (KS2F/R) has been shown to amplify *cis*-AT, *trans*-AT, and hybrid *cis*-AT KSs across diverse bacterial phyla (22–30). When comparing the forward primer (KS2F) to our metagenome-extracted KS domains, we found it aligned best with the *cis*-AT/iterative soil-dominant (n=905) and *cis*-AT/iterative other (n=2257) clades, as >80% of the sequences matched all amino acid residues of the primer (Fig. S12). However, only 46.0% and 40.0% of the sequences in the hybrid *cis*-AT (n=1746) and *trans*-AT (n=831) clades, respectively matched the third amino acid residue (glutamine) from the 3’ end of the KSF primer (Fig. S12). The KS2R reverse primer matched best with the *cis*-AT/iterative soil-dominant clade, as >90% of the sequences matched all amino acid residues. However, sequences from the other clades matched poorly with the 3’ amino acid residue (valine), with only 63.2% of the hybrid *cis*-AT clade, 38.2% of the *trans*-AT clade, and 64.3% of the other *cis*-AT/iterative KS clade matching all amino acid residues.

To date, enediyne KSs have not been reported and PUFA KSs have rarely been reported when the KS2F/R primer set is used (27). This is consistent with our analyses, as <7% of the metagenome-extracted enediyne (n=210) and PUFA (n=1,996) KSs matched the second and fourth amino acid residues (glutamine and arginine/serine, respectively) of the forward primer (Fig. S12). We also evaluated the PUFA-specific primer set pfaA, which was designed based on known *pfaA* KS sequences from GenBank and JGI IMG and used to amplify *pfaA* KSs from marine sediment and seawater samples (22). The 1,170 metagenome-extracted PUFA KS domains matched well with the reverse pfaA primer, with the four amino acid residues (glutamic acid, alanine, histidine, and glycine) closest to the 3’ end matching at 99.7%, 69.7%, 99.0%, and 95.8%, respectively (Fig. S13). However, the metagenome-extracted sequences matched poorly with the forward primer, with 3 of the 4 residues closest to the 3’ end matching at less than 50% (Fig. S13).

Finally, given that PCR generated KS amplicons are likely to be shorter than full length sequences, we asked how this might affect the NaPDoS2 output by shortening the metagenome-extracted *cis*-AT/iterative (n=3,162), hybrid *cis*-AT (n=1,746) and *trans*-AT (n=831) KS domains from full-length (~420 amino acids) to next generation amplicon length (~138 amino acids) and re-analyzing them. Overall, 94.6% of the shortened sequences yielded the same NaPDoS2 classification as their full-length counterparts (95.8% for *cis*-AT/iterative, 94.6% for hybrid *cis*-AT and 90.5% for *trans*-AT). Furthermore, a SSN of the amplicon-length KS domains showed the same clustering pattern as observed in the full-length KS domain SSN (Fig. S4B).

## Discussion

NaPDoS2 provides a rapid method to assess biosynthetic diversity in complex datasets by using KS and C domains to make broader predictions about PKS and NRPS genes and their small molecule products. Using the latest update to this publicly available webtool (15), which features an expanded KS classification scheme and greater capacity for large datasets, we performed an in-depth assessment of type I PKS diversity and distribution in 240 Gbp of metagenomic data derived from eight environmental biomes. We observed significant differences in KS domain composition driven by PUFA KSs in marine biomes and *cis*-AT and hybrid *cis*-AT KSs in soil biomes (Fig. 1a). PUFAs have been suggested to aid in homeoviscous adaptation (22), which could explain why these PKSs are enriched in marine samples. Our analyses also showed that samples from similar biomes shared similar KS diversity, which could reflect biogeographical patterns among KS-containing microbes or environmental selection based on the functional roles of the products they encode. While a recent study of MAGs found no clear skew in relative BGC family content across Earth’s microbiomes (37), we detected biome-specific variations when focusing on diversity within type I KSs, indicating that broad surveys can obscure more subtle but potentially important environmental differences in gene content. Furthermore, we found that only a small subset (<3% on average) of the type I KS domains occurred within MAGs, highlighting the value of using KS sequences to assess PKS diversity. While the drivers of these environmental differences cannot be distinguished here, the KS diversity discerned among biomes can inform natural product discovery efforts and provide insights into the ecological roles of microbial natural products.

The majority of full-length, type I KS domains were classified as *cis*-AT with no further subclassification (Table S2), which aligns with previous genomic explorations of type I KS diversity (42). The search for biome-specific clades within the *cis*-AT/iterative group revealed a soil-dominant clade that mapped almost exclusively to Actinobacteria and grouped with *cis*-AT KS domains associated with experimentally characterized assembly-line PKSs (Fig. 3a). While experimental characterization is biased towards certain taxa, it is possible that certain Actinobacteria with large genomes are uniquely suited for assembly-line megasynthases and that their polyketide products provide a competitive advantage in soil microbiomes. Notably, these results are consistent with previous studies that have shown soil communities to be enriched in Actinobacteria compared to other biomes (43–44). Also aligning with previous work (29–30), our KS phylogeny revealed a sponge-specific clade that was distinct from all KS domains in the MIBiG database (Fig. 3a), thus illustrating the continued potential for natural product discovery from sponge microbiomes. A monomodular clade that was distinct from functionally characterized sequences was also detected among the sequences classified as *cis*-AT/iterative (Figs. 3a, S5). The affiliation of these sequences with Verrucomicrobia, Planctomycetes, and Proteobacteria complements previous studies in *Streptomyces* (16) and suggests that monomodular PKSs are more widely distributed than previously recognized.

Enediyne natural products are rare and of considerable importance as anticancer drugs due to their potent cytotoxicity (40). Our analyses revealed that enediyne KSs were enriched in marine sediments relative to other biomes (Fig. 1b). A phylogeny generated from full-length KS sequences revealed affiliations with diverse phyla such as Proteobacteria, Cyanobacteria, Firmicutes, Bacteroidetes, Chloroflexi, and Spirochaetes (Fig. 4), all taxa from which this class of compounds has yet to be reported. The large marine sediment enediyne lineage is most closely related to a KS domain identified in a Deltaproteobacteria MAG. Searching for enediyne compounds from this taxon could yield new structural diversity in this biomedically important compound class. Additionally, we note the first report of potential enediyne PKSs in anaerobes based on the analysis of NCBI RefSeq genomes. The phylogeny also reveals a soil-specific lineage that mapped exclusively to Actinobacteria and included all the MIBiG-derived enediyne KS domains (Fig. 4). This agrees with previous work showing that Actinobacteria account for most of the enediyne natural products described to date (40–41).

We noticed several patterns in the taxonomic distribution KSs among bacteria. Actinobacteria were the most common taxonomic match for *cis*-AT (44%) and enediyne (28%) KS domains, whereas hybrid *cis*-AT, *trans*-AT, and PUFA domains mostly mapped to Cyanobacteria (24%), Gammaproteobacteria (28%), and Deltaproteobacteria (27%), respectively (Figs. 3b, S6-9). In addition, 23% of the *trans*-AT KS domains mapped to Firmicutes while <4% of the other PKS classes mapped to this phylum. This tracks with previous reports of *trans*-AT PKSs mostly occurring in the Firmicutes and Proteobacteria phyla (45). While microbial gene databases are biased towards cultured representatives, these results suggest that bacterial phyla are differentially enriched in the types of polyketide genes they carry. Noting that 51.1% of the metagenome-extracted KS domains shared a sequence identity of less than 75% with the closest NCBI match (Fig. S11), these taxonomic assignments can be considered tentative and hint at the potential for natural product discovery from poorly studied taxa.

Our analyses revealed that soils and freshwater sediments held greater KS richness than marine sediments and seawater, which contrasts with previous KS amplicon work that reported marine sediments to have greater KS richness than soils (27). While the optimal clustering thresholds to group KS amplicons into meaningful biosynthetic units remain unknown, the KS richness trends we observed in the metagenomes were consistent across thresholds from 70-95%. We also showed that when reducing full-length KS domains to next generation amplicon lengths, 94.6% maintained the same classification (Fig. S4b), thus supporting the use of KS amplicons obtained using next generation sequencing as proxies for full-length type I KS domains.

While metagenomes are not biased towards any specific gene, they are limited in the coverage that can be obtained in complex communities. Conversely, the targeted nature of PCR can capture a more complete picture of KS sequence diversity within a given sample, while being limited to the diversity that can be amplified by the primers. While early studies using the KS2F/R primer set revealed that soil, sponge, and sediment biomes contained significant KS diversity (22–30), a recent study found these primers insufficient for 18 unclassified PKS pathways found in underexplored phyla (31). Capitalizing on our metagenome-derived KS dataset, we found that the KS2F/R primer set aligned best with sequences in the soil-dominant clade within the *cis*-AT/iterative group (Fig. S12). While these primers can recover some hybrid *cis*-AT and *trans*-AT KS sequences, their efficiency for these subclasses could be improved with further primer design. Furthermore, the KS2F/R primer set matches poorly with both PUFA and enediyne KS domains (Fig. S12), as did a PUFA-specific primer set (22) (Fig. S13), suggesting the need for primer modifications that maximize sequence detection within these subclasses.

## Conclusion

An analysis of KS domains in metagenomic data sets using NaPDoS2 revealed linkages between biosynthetic potential and environmental biomes. Through the analysis of 240 Gbps of metagenomic data, we show biome-specific differences in type I KS composition, with PUFA KSs driving the separation in marine biomes, and *cis*-AT and hybrid *cis*-AT KS domains driving the separation in soils. Furthermore, we show that similar biomes share more KS diversity than dissimilar biomes. Phylogenetic analyses of our metagenome-extracted KS domains revealed PTM and enediyne clades that remain unexplored in terms of natural product discovery. Finally, our work revealed that the commonly used KS2F/R primer set is biased towards modular *cis*-AT KS domains and is not well designed to amplify iterative *cis*-AT enediyne and PUFA KSs. Overall, our work highlights the application of KS sequence tags to assess PKS diversity and distributions within large metagenomic datasets.

## Methods

### KS domain identification

One hundred and thirty-seven shotgun metagenomes representing eight biomes (agricultural/forest soil, rhizosphere, peat soil, marine sediment, freshwater sediment, marine water, host-associated, and freshwater) were selected from the JGI IMG database (7) and filtered to exclude contigs <600 nucleotides using a custom script (https://github.com/spodell/NaPDoS2_website/data_management_scripts/size_limit_seqs.pl) resulting in a combined size of 240 Gbp. NaPDoS2 (15) was used to identify type I KS domains using a minimum match length of 200 amino acids and a minimum E-value of 10^-30^. NaPDoS2 KS classifications were verified for a subset of metagenomic sequences across the range of KS domain classes by running the associated contigs through antiSMASH (4) and comparing the output. Additionally, all type I KS domains from the MIBiG database (version 2.0) (6) and associated with MAGS and listed on the JGI IMG database from the 137 metagenomes were also extracted using NaPDoS2 using the same protocols.

### Type I KS domain classification and abundance

KS domains associated with type I PKSs and FASs were identified by NaPDoS2 and further classified as modular *cis*-AT, cis-loading module, olefin synthase, iterative aromatic, iterative PTM, *trans*-AT, hybrid *trans*-AT, hybrid *cis*-AT, PUFA, enediyne, or FAS based on their top matches to the webtool internal database. The relative abundance of each KS domain type was calculated for each metagenome as the number of KS domains/GBP of metagenomic data and compared using a one-way ANOVA followed by Tukey’s HSD test. The total counts for each KS type were rarified to 100 sequences per metagenome using the average from 1000 permutations and transformed into a Bray-Curtis dissimilarity matrix using the Vegan R program (46). This matrix was used to perform a PCoA with significant differences between biomes identified using a permutational ANOVA with the Vegan R program.

### Full-length KS domain analyses

Full-length KS domains were filtered from the total type I metagenomic KS pool and combined with sequences extracted from the MIBiG reference database (6). Metagenomic sequences were considered full-length if they spanned the start residues IAIVG and end residues GTNAH (with some degeneracy at these positions) observed in all type I KS domains within the NaPDoS2 reference database with Geneious ver. 2020.2 (54) used to generate the alignments. A SSN of the full-length sequences was constructed using EFI (47) with an E-value edge calculation of 100 and visualized using Cytoscape (48). Phylogenetic trees were constructed individually for the *trans*-AT, hybrid *cis*-AT, iterative PUFA, iterative enediyne, and *cis*-AT/iterative KS groups identified in the SSN using FastME on the ngphylogeny.fr website (49) with default settings and visualized using ITOL (50).

Due to the large number of KS *cis*-AT/iterative domains identified (3,162), they were grouped by biome and clustered into 70% OBUs using Uclust (51). *Cis*-AT/iterative group KS sequences from the MIBiG database (n=2,149) were similarly clustered into 70% OBUs. FastME trees were constructed using centroid representatives from each OBU with the ngphylogeny.fr webtool (49) under default parameters and visualized using ITOL version 6 (50). No clustering was needed prior to generating the enediyne (n=210), *trans*-AT (n=831), hybrid *cis*-AT (n=1,746) or PUFA (n=1,996) phylogenies.

The genomic context of select KS domains within an uncharacterized clade in the *cis*-AT/iterative phylogeny was visualized by analyzing the relevant metagenomic contigs using antiSMASH 5.0 (4). KS sequences were extracted from the NCBI RefSeq (52) protein database (release number 200) using Blastp ver. 2.11.0 (53) and clustered into 70% OBUs with the metagenome-extracted KS domains. RefSeq genomes that contained KS domains within the same 70% OBU as the uncharacterized clade in the *cis*-AT/iterative phylogeny were then analyzed using antiSMASH to identify the relevant BGCs. A multilocus phylogeny was constructed with the metagenome-extracted BGCs, RefSeq genome-extracted BGCs, and the closest related BGCs from the MIBiG reference database using CORASON (54).

### KS richness, shared diversity, and taxonomic assignments

KS richness and diversity comparisons were made using Geneious ver. 2020.2 (54) with 580 (the minimum number in any biome) full-length KS domains from each biome after clustering into OBUs at 95%, 90%, 80%, and 70% sequence identity using Uclust (51). Average KS richness was estimated at each level based on 10 replicate analyses using the Chao1 index. The number of KS domains in OBUs shared between two biomes was estimated using pairwise comparisons between all biome combinations. Taxonomic affiliations were assigned to full-length KS domains based on the phylum of the closest NCBI (52) Blastp ver. 2.11.0 (53) match (based on E-value).

### Evaluation of KS primer sets

The type I KS primer set KS2F/R (22–30) was aligned with the metagenome-extracted type I KS domains and the percent matching at each amino acid residue calculated using Geneious ver. 2020.2 (55). The PUFA pfaA-specific primer set (22) was similarly analyzed using the metagenome-extracted KS domains that were categorized as PUFA KS01 (or pfaA KS domains) using NaPDoS2.

## Acknowledgements

This work was supported by the National Science Foundation Graduate Research Fellowship Program under Grant no. DGE-2038238 to HWS and the National Institutes of Health grant no. R01GM085770 to PJ. Any opinions, findings, and conclusions or recommendations expressed in this material are those of the author(s) and do not necessarily reflect the views of the National Science Foundation.

## Notes

### Competing Interest Statement

The authors have declared no competing interest.

### Summary of Updates

Addition of supplemental material

## Citations

1. Abdel-Razek AS, El-Naggar ME, Allam A, Morsy OM, Othman SI. 2020. Microbial natural products in drug discovery. Processes 8:470.

2. Pye CR, Bertin MJ, Lokey RS, Gerwick WH, Linington RG. 2017. Retrospective analysis of natural products provides insights for future discovery trends. Proceedings of the National Academy of Sciences 114:5601–5606.

3. Ziemert N, Alanjary M, Weber T. 2016. The evolution of genome mining in microbes – a review. Natural Product Reports 33:988–1005.

4. Blin K, Shaw S, Steinke K, Villebro R, Ziemert N, Lee SY, Medema MH, Weber T. 2019. antiSMASH 5.0: updates to the secondary metabolite genome mining pipeline. Nucleic Acids Research 47:W81–W87.

5. Skinnider MA, Merwin NJ, Johnston CW, Magarvey NA. 2017. PRISM 3: expanded prediction of natural product chemical structures from microbial genomes. Nucleic Acids Research 45:W49–W54.

6. Kautsar SA, Blin K, Shaw S, Navarro-Muñoz JC, Terlouw BR, van der Hooft JJJ, van Santen JA, Tracanna V, Duran HGS, Andreu VP, Selem-Mojica N, Alanjary M, Robinson SL, Lund G, Epstein SC, Sisto AC, Charkoudian LK, Collemare J, Linington RG, Weber T, Medema MH. 2020. MIBiG 2.0: a repository for biosynthetic gene clusters of known function. Nucleic Acids Research 48:D454–D458.

7. Palaniappan K, Chen IMA, Chu K, Ratner A, Seshadri R, Kyrpides NC, Ivanova NN, Mouncey NJ. 2020. IMG-ABC v. 5.0: an update to the IMG/Atlas of biosynthetic gene clusters knowledgebase. Nucleic Acids Research. 48:D422–D430.

8. Nivina A, Yuet KP, Hsu J, Khosla C. 2019. Evolution and Diversity of Assembly-Line Polyketide Synthases: Focus Review. Chemical Reviews 119:12524–12547.

9. Weissman KJ. 2004. Polyketide biosynthesis: understanding and exploiting modularity. Philosophical Transactions of the Royal Society of London Series A, Mathematical, Physical and Engineering Sciences 362:2671–2690.

10. Fischbach MA, Walsh CT. 2006. Assembly-line enzymology for polyketide and nonribosomal peptide antibiotics: logic, machinery, and mechanisms. Chemical Reviews 106:3468–3496.

11. Shen B. 2003. Polyketide biosynthesis beyond the type I, II and III polyketide synthase paradigms. Current Opinion in Chemical Biology 7:285–295.

12. Chen H, Du L. 2016. Iterative polyketide biosynthesis by modular polyketide synthases in bacteria. Applied Microbiology and Biotechnology 100:541–557.

13. Piel J. 2010. Biosynthesis of polyketides by trans-AT polyketide synthases. Natural Product Reports 27:996–1047.

14. Ziemert N, Podell S, Penn K, Badger JH, Allen E, Jensen PR. 2012. The natural product domain seeker NaPDoS: A phylogeny based bioinformatic tool to classify secondary metabolite gene diversity. PLoS One 7:e34064.

15. Klau LJ, Podell S, Creamer KE, Demko AM, Singh HWS, Allen EE, Moore BS, Ziemert N, Letzel AC, Jensen PR. 2022. The Natural Product Domain Seeker version 2 (NaPDoS2) webtool relates ketosynthase phylogeny to biosynthetic function. Journal of Biological Chemistry 298:102480.

16. Wang B, Guo F, Huang C, Zhao H. 2020. Unraveling the iterative type I polyketide synthases hidden in Streptomyces. Proceedings of the National Academy of Sciences 117:8449–8454.

17. Miyanaga A, Kudo F, Eguchi T. 2018. Protein–protein interactions in polyketide synthase– nonribosomal peptide synthetase hybrid assembly lines. Natural Product Reports 35:1185–1209.

18. Moffitt MC, Neilan BA. 2003. Evolutionary affiliations within the superfamily of ketosynthases reflect complex pathway associations. Journal of Molecular Evolution 56:446–457.

19. Ayuso-Sacido A, Genilloud O. 2005. New PCR Primers for the Screening of NRPS and PKS-I Systems in Actinomycetes: Detection and Distribution of These Biosynthetic Gene Sequences in Major Taxonomic Groups. Microbial Ecology 49:10–24.

20. Wawrik B, Kerkhof L, Zylstra GJ, Kukor JJ. 2005. Identification of Unique Type II Polyketide Synthase Genes in Soil. Applied and Environmental Microbiology 71:2232–2238.

21. Rascher A, Hu Z, Viswanathan N, Schirmer A, Reid R, Nierman WC, Lewis M, Hutchinson CR. 2003. Cloning and characterization of a gene cluster for geldanamycin production in Streptomyces hygroscopicus NRRL 3602. FEMS Microbiology Letters 218:223–230.

22. Shulse CN, Allen EE. 2011. Diversity and distribution of microbial long-chain fatty acid biosynthetic genes in the marine environment. Environmental Microbiology 13:684–695.

23. Charlop-Powers Z, Owen JG, Reddy BVB, Ternei MA, Brady SF. 2014. Chemical-biogeographic survey of secondary metabolism in soil. Proceedings of the National Academy of Sciences 111:3757–3762.

24. Charlop-Powers Z, Owen JG, Reddy BVB, Ternei MA, Guimarães DO, de Frias UA, Pupo MT, Seepe P, Feng Z, Brady SF. 2015. Global biogeographic sampling of bacterial secondary metabolism. eLife 4:e05048.

25. Libis V, Antonovsky N, Zhang M, Shang Z, Montiel D, Maniko J, Ternei MA, Calle PY, Lemetre C, Owen JG, Brady SF. 2019. Uncovering the biosynthetic potential of rare metagenomic DNA using co-occurrence network analysis of targeted sequences. Nature Communications 10:3848.

26. Bech PK, Lysdal KL, Gram L, Bentzon-Tilia M, Strube ML. 2020. Marine sediments hold an untapped potential for novel taxonomic and bioactive bacterial diversity. Msystems 5:e00782–20.

27. Rego A, Sousa AGG, Santos JP, Pascoal F, Canário J, Leão PN, Magalhães C. 2020. Diversity of bacterial biosynthetic genes in maritime antarctica. Microorganisms 8:279.

28. Elfeki M, Mantri S, Clark CM, Green SJ, Ziemert N, Murphy BT. 2021. Evaluating the distribution of bacterial natural product biosynthetic genes across Lake Huron sediment. ACS Chemical Biology 16:2623–2631.

29. Hochmuth T, Piel J. 2009. Polyketide synthases of bacterial symbionts in sponges – Evolution-based applications in natural products research. Phytochemistry 70:1841–1849.

30. Sala DG, Hochmuth T, Teta R, Costantino V, Mangoni A. 2014. Polyketide synthases in the microbiome of the marine sponge *Plakortis halichondrioides:* A metagenomic update. Marine Drugs 12:5425–5440.

31. Crits-Christoph A, Diamond S, Butterfield CN, Thomas BC, Banfield JF. 2018. Novel soil bacteria possess diverse genes for secondary metabolite biosynthesis. Nature 558:440–444.

32. Paoli L, Ruscheweyh HJ, Forneris CC, Hubrich F, Kautsar S, Bhushan A, Lotti A, Clayssen Q, Salazar G, Milanese A, Carlstrom CI, Papadopoulou C, Gehrig D, Karasikov M, Mustafa H, Larralde M, Carroll LM, Sánchez P, Zayed AA, Cronin DR, Acinas SG, Bork P, Bowler C, Delmont TO, Gasol JM, Gossert AD, Kahles L, Sullivan MB, Wincker P, Zeller G, Robinson SL, Piel J, Sunagawa S. 2022. Biosynthetic potential of the global ocean microbiome. Nature 607:111–118.

33. Parks DH, Rinke C, Chuvochina M, Chaumeil PA, Woodcroft BJ, Evans PN, Hugenholtz P, Tyson GW. 2017. Recovery of nearly 8,000 metagenome-assembled genomes substantially expands the tree of life. Nature Microbiology 2:1533–1542.

34. Sugimoto Y, Camacho FR, Wang S, Chankhamjon P, Odabas A, Biswas A, Jeffrey PD, Donia MS. 2019. A metagenomic strategy for harnessing the chemical repertoire of the human microbiome. Science 366:eaax9176.

35. Carrión VJ, Perez-Jaramillo J, Cordovez V, Tracanna V, Hollander M, Ruiz-Buck D, Mendes LW, van Ijcken WFJ, Gomez-Exposito R, Elsayed SS, Mohanraju P, Arifah A, Oost J, Paulson JN, Mendes R, van Wezel GP, Medema MH, Raaijmakers JM. 2019. Pathogen-induced activation of disease-suppressive functions in the endophytic root microbiome. Science 366:606–612.

36. Storey MA, Andreassend SK, Bracegirdle J, Brown A, Keyzers RA, Ackerley DF, Northcote PT, and Owen JG. 2020. Metagenomic exploration of the marine sponge *Mycale hentscheli* uncovers multiple polyketide-producing bacterial symbionts. mBio 11:e02997–19.

37. Nayfach S, Roux S, Seshadri R, Udwary D, Varghese N, Schulz F, Wu D, Paez-Espino D, Chen IM, Huntemann M. 2020. A genomic catalog of Earth’s microbiomes. Nature Biotechnology 39:499–509.

38. Ziemert N, Lechner A, Wietz M, Jensen PR. 2014. Diversity and evolution of secondary metabolism in the marine actinomycete genus *Salinispora*. Proceedings of the National Academy of Sciences 111:E1130–E1139.

39. Blodgett JA, Oh VDC, Cao S, Currie CR, Kolter R, Clardy J. 2010. Common biosynthetic origins for polycyclic tetramate macrolactams from phylogenetically diverse bacteria. Proceedings of the National Academy of Sciences 107:11692–11697.

40. Shen B, Hindra, Yan X, Huang T, Ge H, Yang D, Teng Q, Rudolf JD, Lohman JR. 2015. Enediynes: Exploration of microbial genomics to discover new anticancer drug leads. Bioorganic & Medicinal Chemistry Letters 25:9–15.

41. Rudolf JD, Yan X, Shen B. 2016. Genome neighborhood network reveals insights into enediyne biosynthesis and facilitates prediction and prioritization for discovery. Journal of Industrial Microbiology and Biotechnology 43:261–276.

42. O’Brien RV, Davis RW, Khosla C, Hillenmeyer ME. 2014. Computational identification and analysis of orphan assembly-line polyketide synthases. The Journal of Antibiotics 67:89–97.

43. Delgado-Baquerizo M, Oliverio AM, Brewer TE, Benavent-González A, Eldridge DJ, Bardgett RD, Maestre FT, Singh BK, Fierer N. 2018. A global atlas of the dominant bacteria found in soil. Science 359:320–325.

44. Hoshino T, Doi H, Uramoto G-I, Wörmer L, Adhikari RR, Xiao N, Morono Y, D’Hondt S, Hinrichs K-U, Inagaki F. 2020. Global diversity of microbial communities in marine sediment. Proceedings of the National Academy of Sciences 117:27587–27597.

45. Nguyen T, Ishida K, Jenke-Kodama H, Dittmann E, Gurgui C, Hochmuth T, Taudien S, Platzer M, Hertweck C, Piel J. 2008. Exploiting the mosaic structure of trans-acyltransferase polyketide synthases for natural product discovery and pathway dissection. Nature Biotechnology 26:225–233.

46. Oksanen J, Blanchet FG, Friendly M, Kindt R, Legendre P, McGlinn D, Minchin PR, O’Hara RB, Simpson GL, Solymos P, Henry M, Stevens H, Szoecs E, Wagner H. 2020. vegan: Community Ecology Package. R package version 2.5-7.

47. Zallot R, Oberg N, Gerlt JA. 2019. The EFI web resource for genomic enzymology tools: Leveraging protein, genome, and metagenome databases to discover novel enzymes and metabolic pathways. Biochemistry 58:4169–4182.

48. Shannon P, Markiel A, Ozier O, Baliga NS, Wang JT, Ramage D, Amin N, Schwikowski B, Ideker T. 2003. Cytoscape: a software environment for integrated models of biomolecular interaction networks. Genome Research 13:2498–2504.

49. Lemoine F, Correia D, Lefort V, Doppelt-Azeroual O, Mareuil F, Cohen-Boulakia S, Gascuel O. 2019. NGPhylogeny.fr: new generation phylogenetic services for non-specialists. Nucleic Acids Research 47:W260–W265.

50. Letunic I, Bork P. 2021. Interactive Tree Of Life (iTOL) v5: an online tool for phylogenetic tree display and annotation. Nucleic Acids Research. 49(W1), W293–W296.

51. Edgar RC. 2010. Search and clustering orders of magnitude faster than BLAST. Bioinformatics 26:2460–2461.

52. O’Leary NA, Wright MW, Brister JR, Ciufo S, Haddad D, McVeigh R, Rajput B, Robbertse B, Smith-White B, Ako-Adjei D, Astashyn A, Badretdin A, Bao Y, Blinkova O, Brover V, Chetvernin V, Choi J, Cox E, Ermolaeva O, Farrell CM, Goldfarb T, Gupta T, Haft D, Hatcher E, Hlavina W, Joardar VS, Kodali VK, Li W, Maglott D, Masterson P, McGarvey KM, Murphy MR, O’Neill K, Pujar S, Rangwala SH, Rausch D, Riddick LD, Schoch C, Shkeda A, Storz SS, Sun H, Thibaud-Nissen F, Tolstoy I, Tully RE, Vatsan AR, Wallin C, Webb D, Wu W, Landrum MJ, Kimchi A, Tatusova T, DiCuccio M, Kitts P, Murphy TD, Pruitt KD. 2016. Reference sequence (RefSeq) database at NCBI: current status, taxonomic expansion, and functional annotation. Nucleic Acids Research 44:D733–D745.

53. Camacho C, Coulouris G, Avagyan V, Ma N, Papadopoulos J, Bealer K, Madden TL. 2009. BLAST+: architecture and applications. BMC Bioinformatics 10:421.

54. Navarro-Muñoz JC, Selem-Mojica N, Mullowney MW, Kautsar SA, Tryon JH, Parkinson EI, De Los Santos ELC, Yeong M, Cruz-Morales P, Abubucker S, Roeters A, Lokhorst W, Fernandez-Guerra A, Cappelini LTD, Goering AW, Thomson RJ, Metcalf WW, Kelleher NL, Barona-Gomez F, Medema MH. 2020. A computational framework to explore large-scale biosynthetic diversity. Nature Chemical Biology 16:60–68.

55. Kearse M, Moir R, Wilson A, Stones-Havas S, Cheung M, Sturrock S, Buxton S, Cooper A, Markowitz S, Duran C, Thierer T, Ashton B, Meintjes P, Drummond A. 2012. Geneious Basic: an integrated and extendable desktop software platform for the organization and analysis of sequence data. Bioinformatics 28:1647–1649.

