## Supplemental figures and tables for "Metagenomic Data Reveal Type I Polyketide Synthase Distributions Across Biomes"

**Table S1 (attached separately) - Summary table of all metagenomes analyzed using NaPDoS2 .** All metagenomes are listed along with the biome type they fall under and the total size of the metagenome (base pairs).

| Class | Subclass | All biomes | Forest / Agricultural soil | Peat Soil | Rhizosphere | Marine Sediment | Freshwater Sediment | Host-associated | Seawater | Freshwater |
|---|---|---|---|---|---|---|---|---|---|---|
| Modular *cis*-AT | No subclass | 13186 | 3381 | 2539 | 1790 | 951 | 1070 | 1275 | 951 | 1229 |
| | Hybrid KS | 6655 | 1452 | 1105 | 1316 | 907 | 687 | 326 | 318 | 544 |
| | Loading module | 864 | 215 | 139 | 152 | 129 | 54 | 98 | 21 | 56 |
| | Olefin synthase | 719 | 135 | 79 | 121 | 98 | 58 | 27 | 99 | 102 |
| Iterative *cis*-AT | PUFA | 7346 | 460 | 791 | 386 | 2750 | 899 | 134 | 1141 | 785 |
| | No subclass | 1523 | 264 | 178 | 218 | 204 | 229 | 104 | 204 | 122 |
| | Enediyne | 1044 | 173 | 132 | 153 | 403 | 66 | 12 | 49 | 56 |
| | Aromatic | 355 | 140 | 48 | 79 | 4 | 34 | 10 | 7 | 33 |
| | PTM-type | 207 | 44 | 39 | 36 | 21 | 14 | 6 | 10 | 37 |
| *trans*-AT | No subclass | 3061 | 683 | 625 | 473 | 345 | 113 | 291 | 177 | 354 |
| | Hybrid KS | 156 | 26 | 35 | 27 | 12 | 10 | 15 | 16 | 15 |
| | Total KSs | 35116 | 6973 | 5710 | 4751 | 5824 | 3234 | 2298 | 2993 | 3333 |
| Type I FAS | | 409 | 93 | 36 | 58 | 10 | 35 | 94 | 45 | 38 |

**Table S2 - Type I KS hits classified by NaPDoS2.** Type I KS domains listed by NaPDoS2 class and subclass across eight biomes, with the total KS hits across all biomes listed in the third column. The total number of Type I KS hits across all classes is listed in the second to last row. Type I FAS hits were not included in these totals and are listed in the last row.

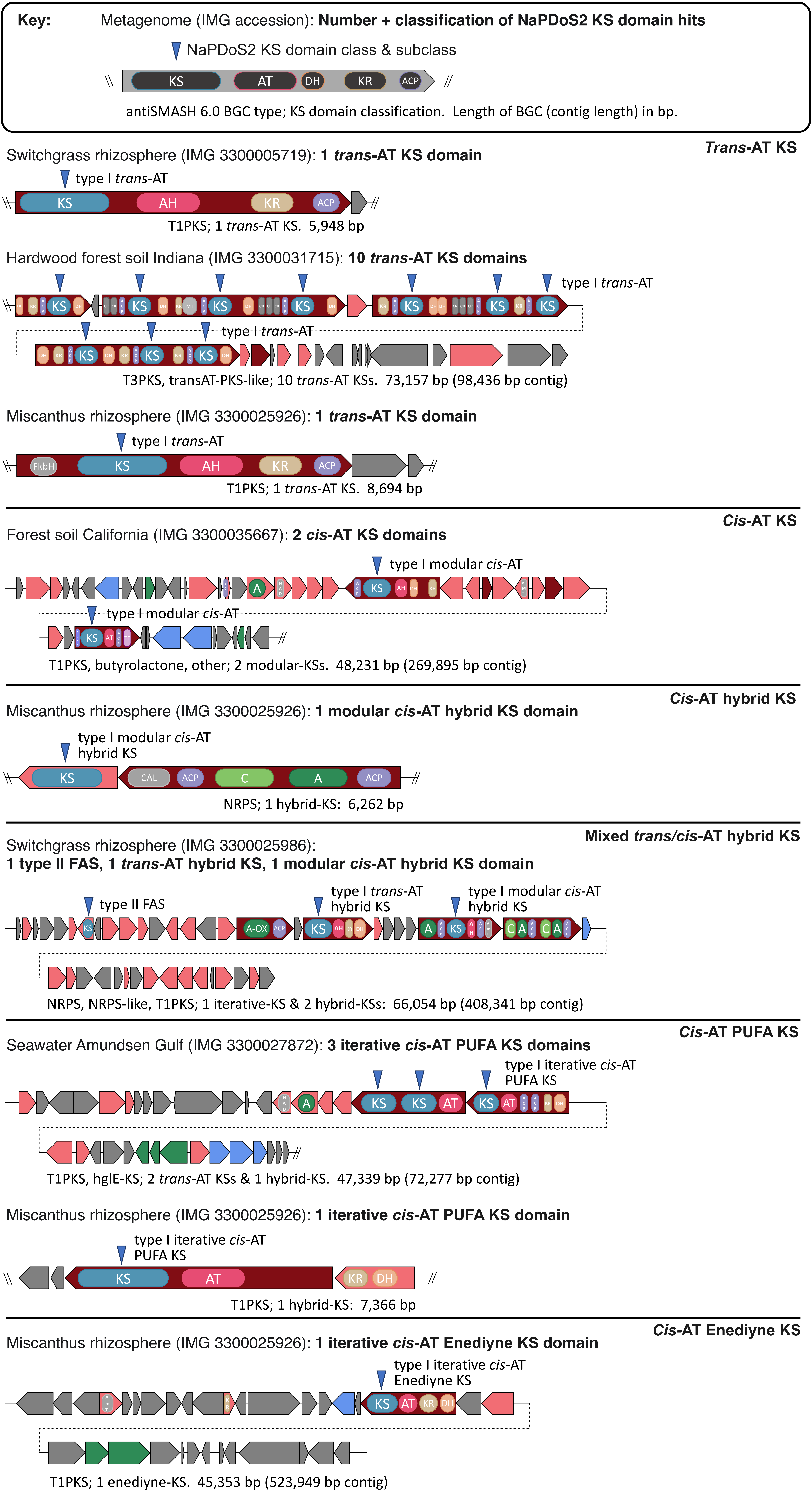
Key: Metagenome (IMG accession): Number + classification of NaPDoS2 KS domain hits
NaPDoS2 KS domain class & subclass
KS
AT
DH
KR
ACP
antiSMASH 6.0 BGC type; KS domain classification. Length of BGC (contig length) in bp.
Trans-AT KS
Switchgrass rhizosphere (IMG 3300005719): 1 trans-AT KS domain
type I trans-AT
KS
AH
KR
ACP
T1PKS; 1 trans-AT KS. 5,948 bp
Hardwood forest soil Indiana (IMG 3300031715): 10 trans-AT KS domains
type I trans-AT
type I trans-AT
T3PKS, transAT-PKS-like; 10 trans-AT KSs. 73,157 bp (98,436 bp contig)
Miscanthus rhizosphere (IMG 3300025926): 1 trans-AT KS domain
type I trans-AT
FkbH
KS
AH
KR
ACP
T1PKS; 1 trans-AT KS. 8,694 bp
Cis-AT KS
Forest soil California (IMG 3300035667): 2 cis-AT KS domains
type I modular cis-AT
type I modular cis-AT
T1PKS, butyrolactone, other; 2 modular-KSs. 48,231 bp (269,895 bp contig)
Cis-AT hybrid KS
Miscanthus rhizosphere (IMG 3300025926): 1 modular cis-AT hybrid KS domain
type I modular cis-AT hybrid KS
KS
CAL
ACP
C
A
ACP
NRPS; 1 hybrid-KS: 6,262 bp
Mixed trans/cis-AT hybrid KS
Switchgrass rhizosphere (IMG 3300025986):
1 type II FAS, 1 trans-AT hybrid KS, 1 modular cis-AT hybrid KS domain
type II FAS
type I trans-AT hybrid KS
type I modular cis-AT hybrid KS
A-OX
ACP
NRPS, NRPS-like, T1PKS; 1 iterative-KS & 2 hybrid-KSs: 66,054 bp (408,341 bp contig)
Cis-AT PUFA KS
Seawater Amundsen Gulf (IMG 3300027872): 3 iterative cis-AT PUFA KS domains
type I iterative cis-AT PUFA KS
KS
KS
AT
KS
AT
T1PKS, hglE-KS; 2 trans-AT KSs & 1 hybrid-KS. 47,339 bp (72,277 bp contig)
Miscanthus rhizosphere (IMG 3300025926): 1 iterative cis-AT PUFA KS domain
type I iterative cis-AT PUFA KS
KS
AT
KR
DH
T1PKS; 1 hybrid-KS: 7,366 bp
Cis-AT Enediyne KS
Miscanthus rhizosphere (IMG 3300025926): 1 iterative cis-AT Enediyne KS domain
type I iterative cis-AT Enediyne KS
KS
AT
KR
DH
T1PKS; 1 enediyne-KS. 45,353 bp (523,949 bp contig)

**Figure S1 - Examples of BGC and gene neighborhood context for NaPDoS2 KS hits.** Random KS hits of various subclasses including trans-AT, cis-AT, cis-AT hybrid, mixed trans/cis-AT hybrid, PUFA, cis-AT PUFA, and cis-AT enediyne) identified by NaPDoS2 were located in the metagenomes they were detected/extracted from by using blastP of the KS domains against the metagenome in the JGI IMG interface. The entire metagenome scaffold/contig with the KS hit was extracted, and run through antiSMASH 6 to identify BGC regions and important biosynthetic genes; transATor, and "PKS/NRPS Analysis Web-site" (http://nrps.igs.umaryland.edu/) for relevant BGC domain detection. The BGCs were drawn and colored as determined by antiSMASH6 (maroon, core biosynthetic gene; pink, additional biosynthetic gene; blue, transport-related genes; green, regulatory genes, gray, other genes); domain position and function were drawn and colored according to antiSMASH, transATor, and NRPS.IGS (blue, KS ketosynthase; pink: AH acyl hydrolase, AT acyl transferase; sand, KR ketoreductase; pale purple, ACP Phosphopantetheine acyl carrier protein; orange, DH dehydratase; dark gray, CR crotonase; light gray: MT methyltransferase, FkbH domain, CAL Co-enzyme A ligase domain, NAD Male sterility protein, oMT Oxygen methyltransferase, AmT aminotransferase; light pink, TE thioesterase; dark blue, dock PKS terminal docking domain; light green, C condensation domain; dark green: A adenylation domain, A-OX Adenylation domain with integrated oxidase). Blue arrows point to KS hits that NaPDoS2 detected and classified from each metagenome shown in the BGC context; arrows are labeled with the NaPDoS2 classification.

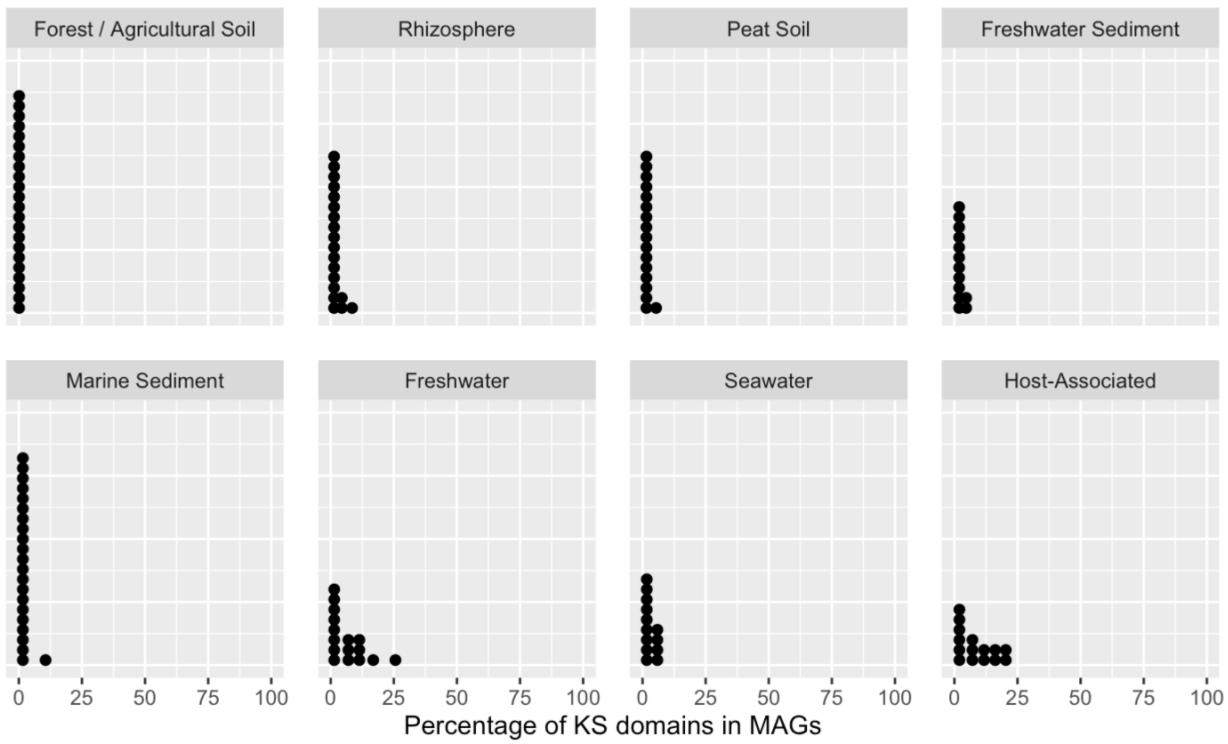


**Figure S2 - Percentage of metagenomic KS domains detected within MAGs.** Individual metagenomes (137 in total from IMG/M) are represented by black circles. MAGs were binned for each metagenome using MetaBAT according to the JGI IMG automated pipeline. The metagenomes and MAGs from those metagenomes were analyzed independently using NaPDoS2, and the percentage of KS domains in MAGs compared to the entire metagenome was plotted.

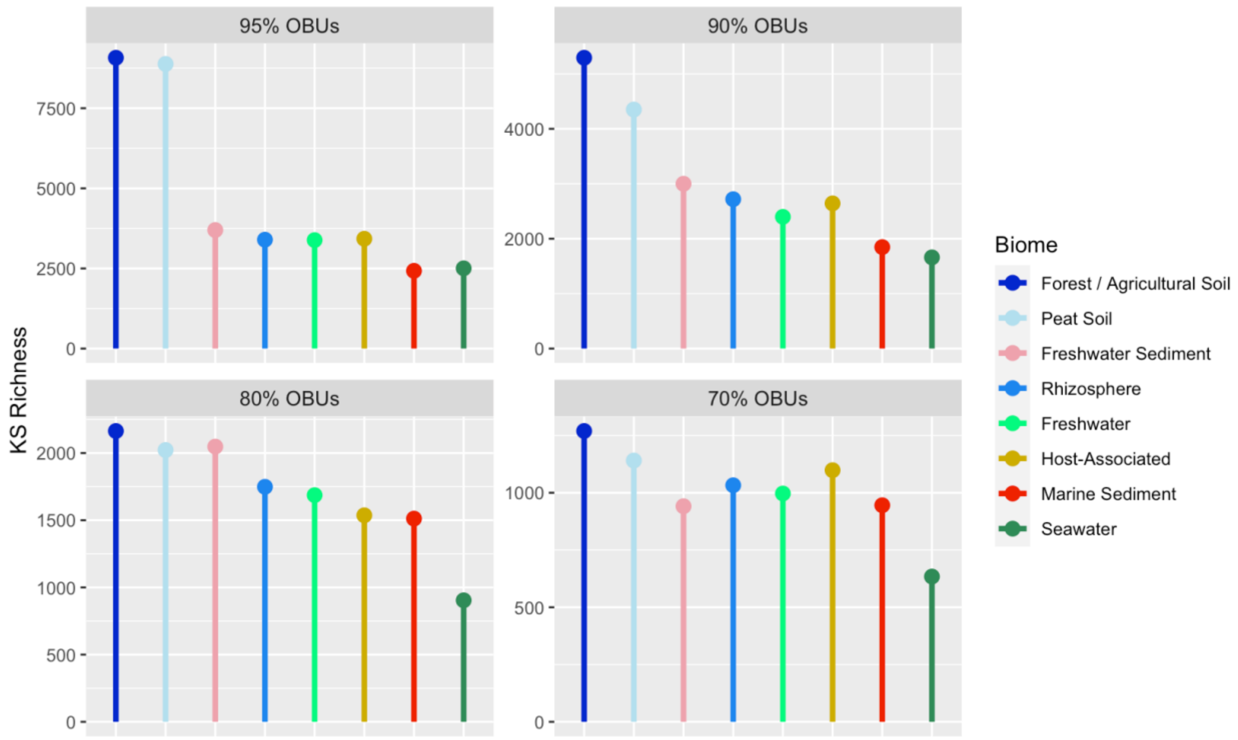


**Figure S3 - KS richness across biomes.** Bar plot showing KS richness, calculated using the Chao1 richness index across eight biomes. For each biome, 581 full-length KS domains were randomly selected and clustered at four different OTU thresholds (panels) ranging from 70% to 95%. The results represent the average of 10 analyses per biome.

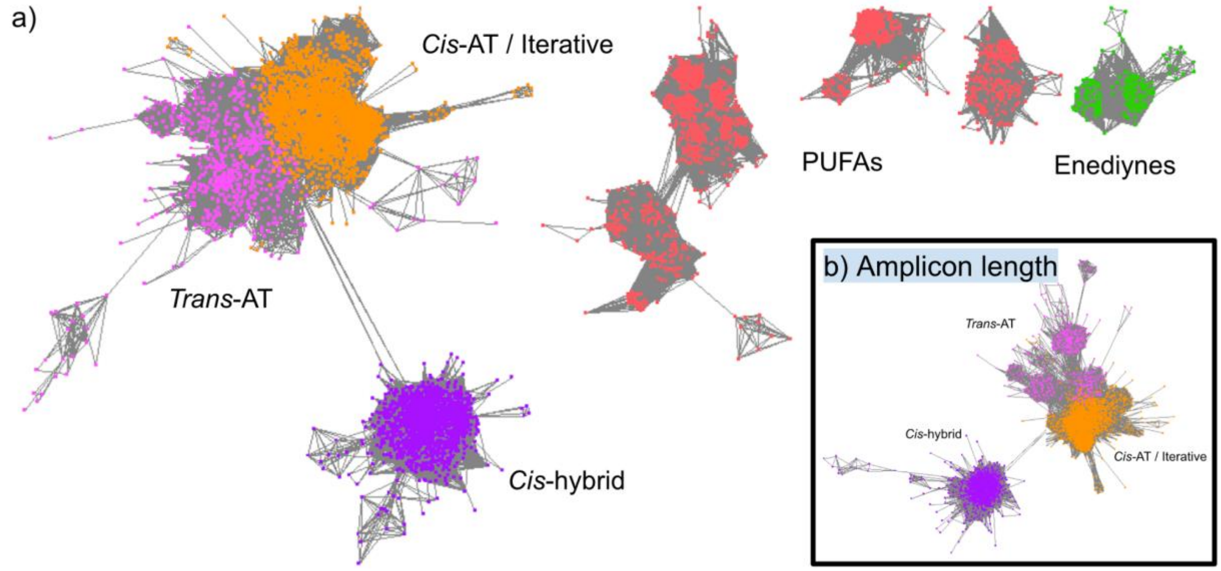


**Figure S4 - Type I KS domains colored by NaPDoS2 classification.** A) An SSN of all full-length metagenome-extracted KS domains (main) was constructed using EFI and visualized using Cytoscape. KS domain sequences (amino acids, average length = 420 aa) were colored according to their NaPDoS2 classification, with five clear groups appearing. KS domains from *cis*-hybrid (purple), trans-AT (pink), PUFAs (red) and enediynes (green) formed four subclass-specific groups. Additionally, KS domains from the *cis*-AT modular, *cis*-loading modules, OLS, iterative aromatics and iterative PTMs classes formed a fifth group that was termed the *cis*-AT/iterative group (orange). B) All KS domains from the *cis*-AT/iterative, trans-AT and cis-hybrid groups were shortened to amplicon lengths (amino acids, average length = 138 aa) and used to create a SSN. The same separation of these three clades was observed at amplicon lengths (inset,B) as for full-length KS domains (main,A).

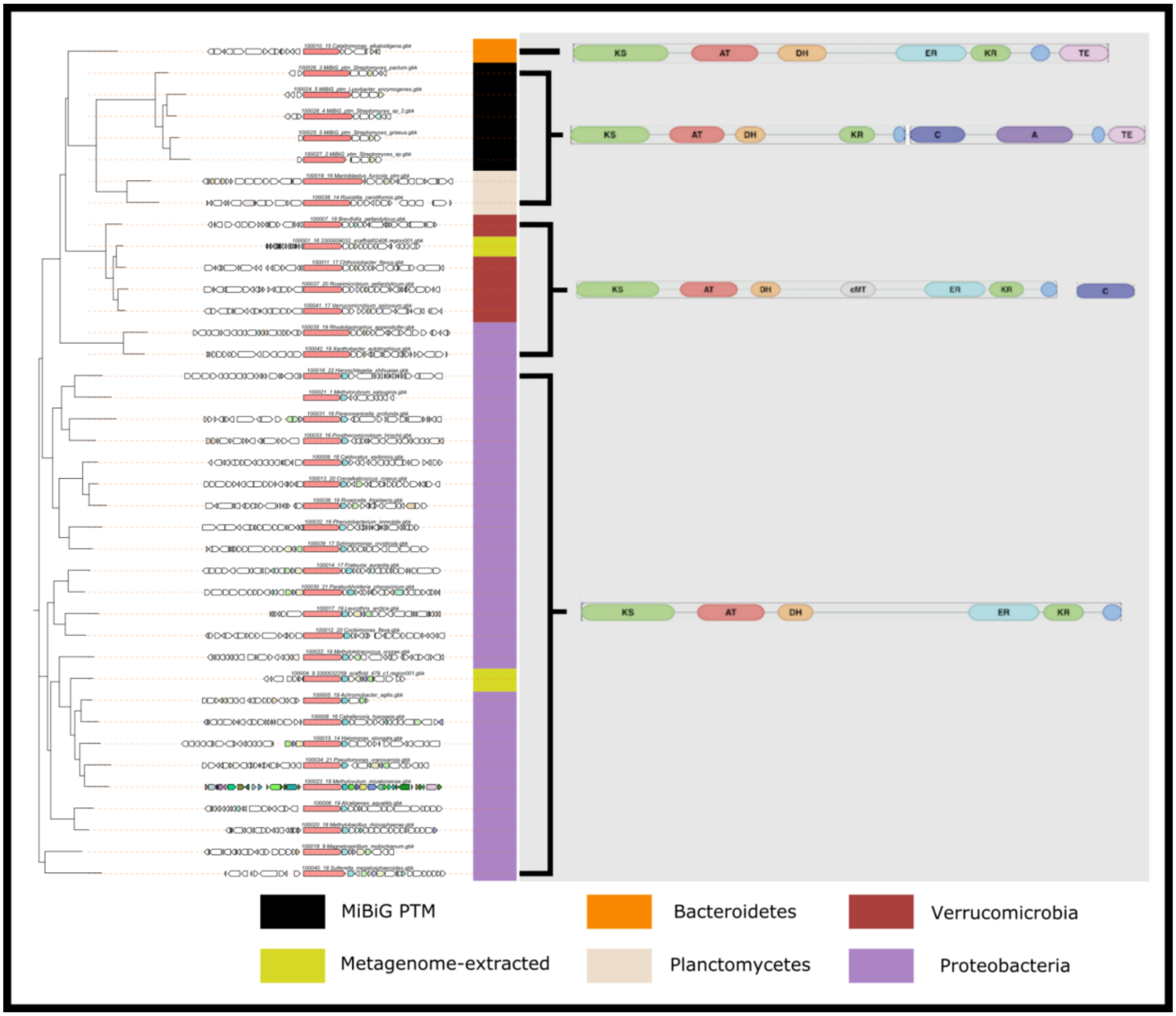


**Figure S5 - Multilocus phylogeny of Monomodular clade adjacent to MIBiG PTMs.** To contextualize a KS clade from the *cis*-AT/iterative phylogeny (Fig. 2a, pink) that was exclusive of KS sequences from MiBIG, a multilocus phylogeny was used. Two full-length BGCs were pulled from metagenomic contigs from this KS clade using antiSMASH (yellow), and BGCs from RefSeq genomes that grouped into KS OBUs (70%) from this clade were also extracted (colored by taxa of RefSeq genome). Included in this phylogeny as reference points are full-length BGCs corresponding to the closest MIBiG representatives to this clade (Iterative PTM BGCs in black). To the right of the phylogeny, the architecture of the polyketide synthase gene as visualized in the AntiSMASH outputs are shown.

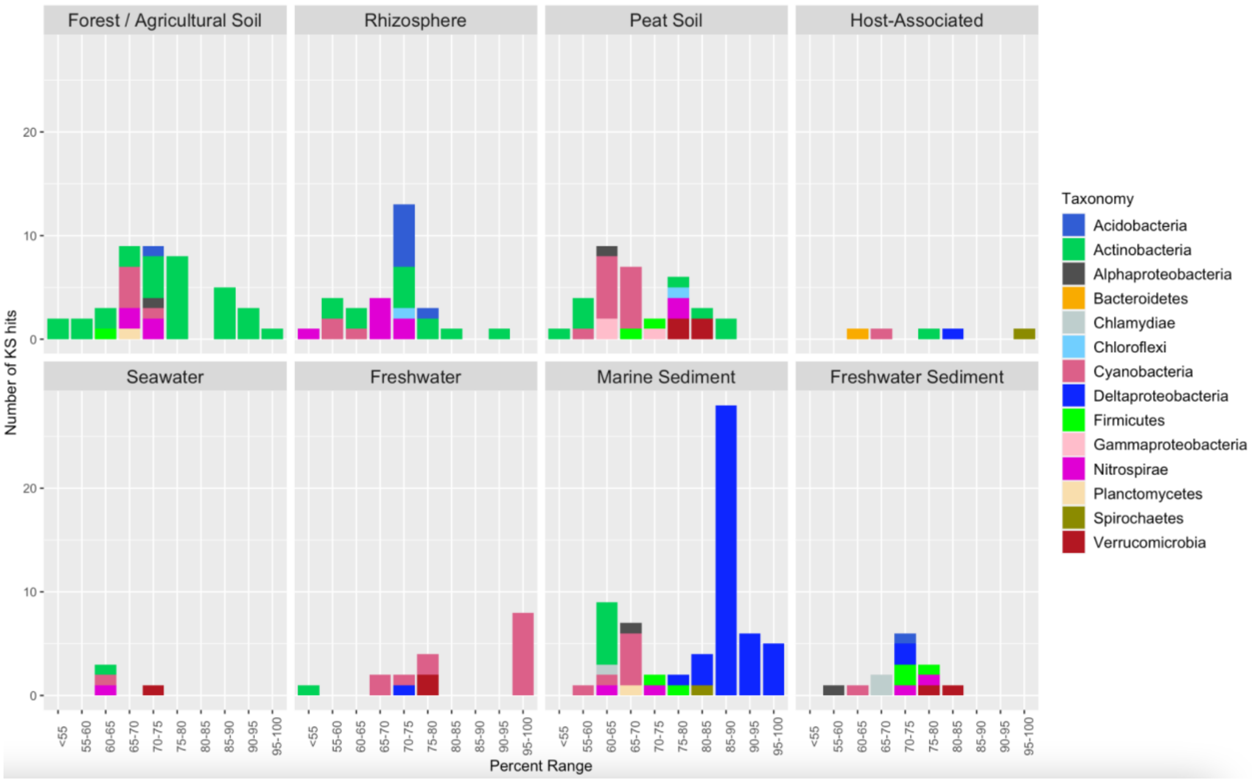


**Figure S6 - Enediyne KS domain distributions across biomes.** Stacked bar charts indicate the phylum-level taxonomic composition and abundance (y-axis) of full-length metagenome-extracted enediyne KS domains for eight biomes grouped based on percent identity with the closest NCBI BLASTp database match (x-axis).

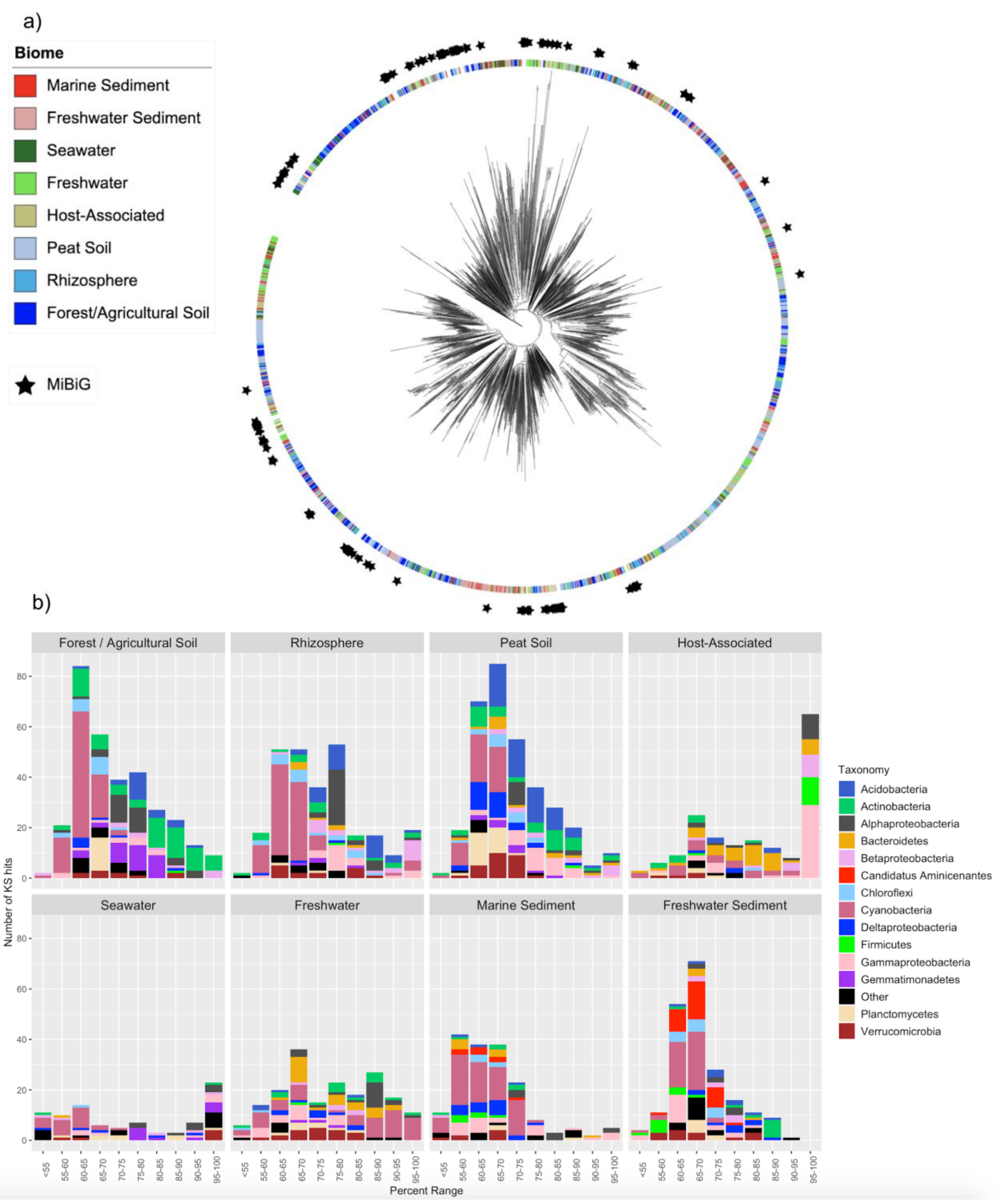


**Figure S7 - *Cis*-hybrid KS domain phylogeny and distributions across biomes.** A) A FastME phylogeny was generated from full-length metagenome-extracted *cis*-hybrid KS domains (n=1746, colored by biome) with the position of MiBiG-extracted *cis*-hybrid KS domains shown as black stars. B) Stacked bar charts indicate the phylum-level taxonomic composition and abundance (y-axis) of full-length metagenome-extracted *cis*-AT KS domains for eight biomes grouped based on percent identity with the closest NCBI BLASTp database match (x-axis).

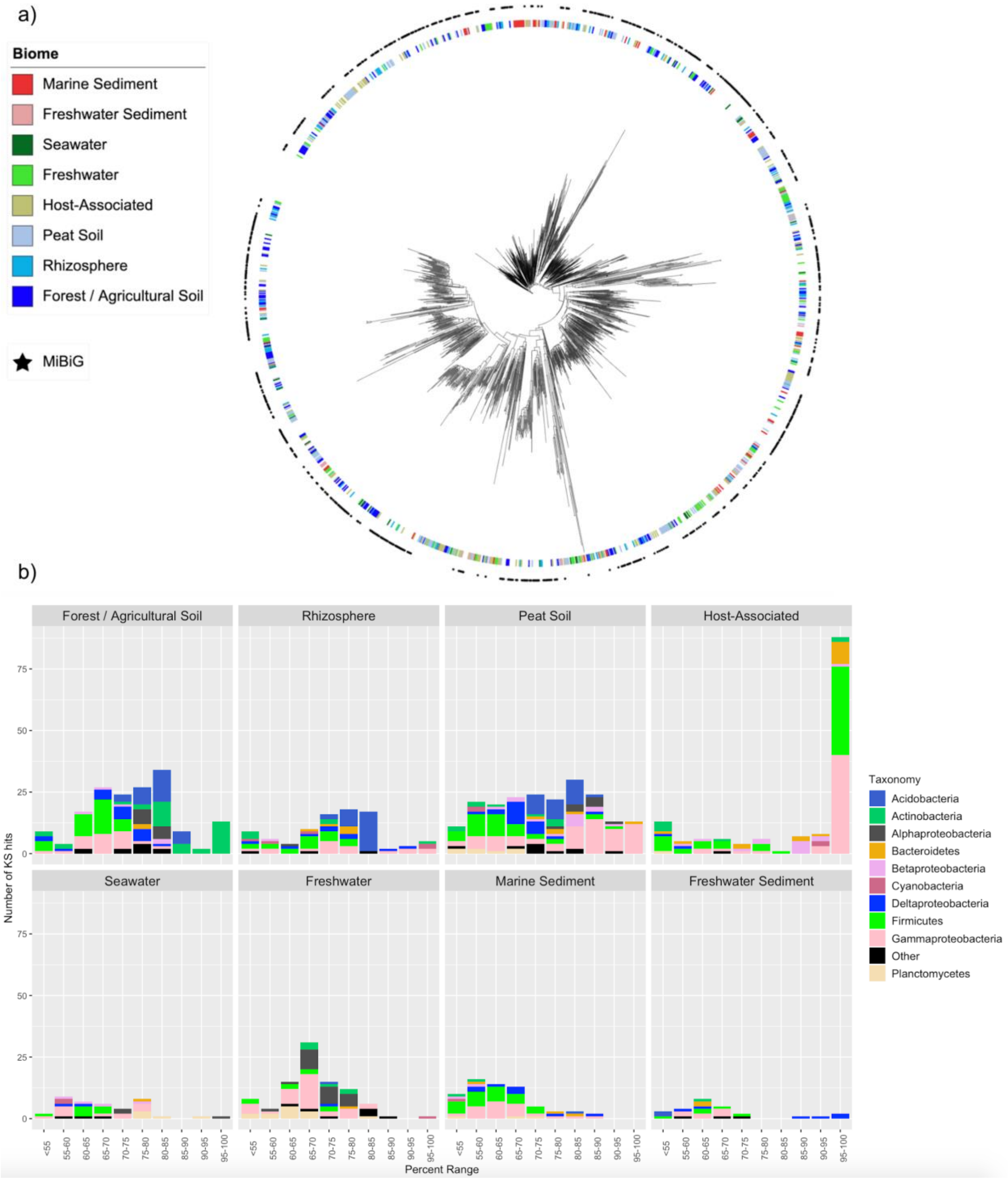


**Figure S8 - *Trans*-AT KSs domain phylogeny and distributions across biomes.** A) A FastME phylogeny generated from all full-length, metagenome-extracted *trans*-AT KS domains (n=831, colored by biome) with the position of MiBiG-extracted *trans*-AT KS domains shown as black stars). B) Stacked bar charts indicate the phylum-level taxonomic composition and abundance (y-axis) of full-length metagenome-extracted *trans*-AT KS domains for eight biomes grouped based on percent identity with the closest NCBI BLASTp database match (x-axis).

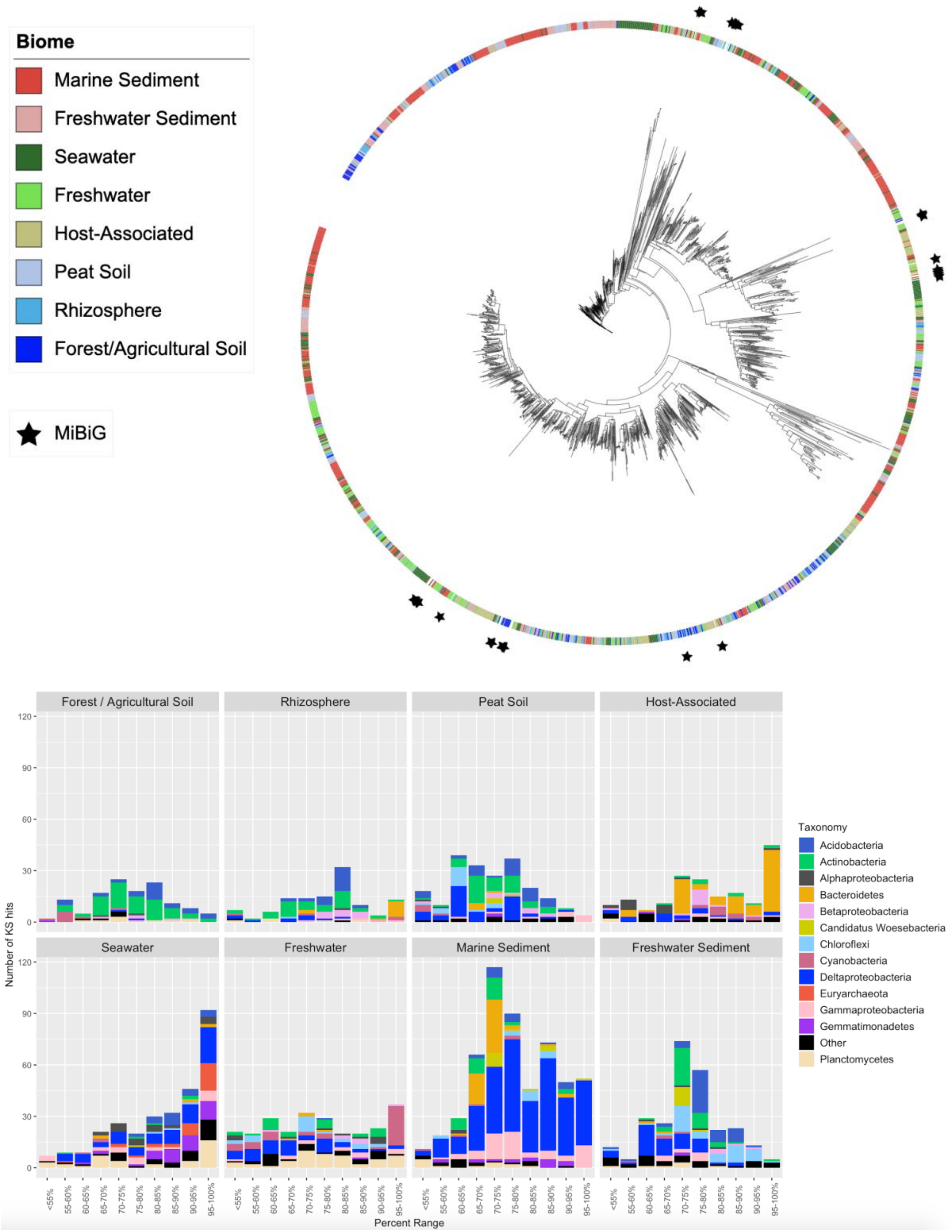


**Figure S9 - PUFA KS domain phylogeny and distributions across biomes.** A) FastME phylogeny generated from full-length metagenome-extracted PUFA KS domains (n=1996, colored by biome) with the position of MIBiG PUFA KS domains shown as black stars. B)

Stacked bar charts indicate the phylum-level taxonomic composition and abundance (y-axis) of full-length metagenome-extracted PUFA KS domains for eight biomes grouped based on percent identity with the closest NCBI BLASTp database match (x-axis).

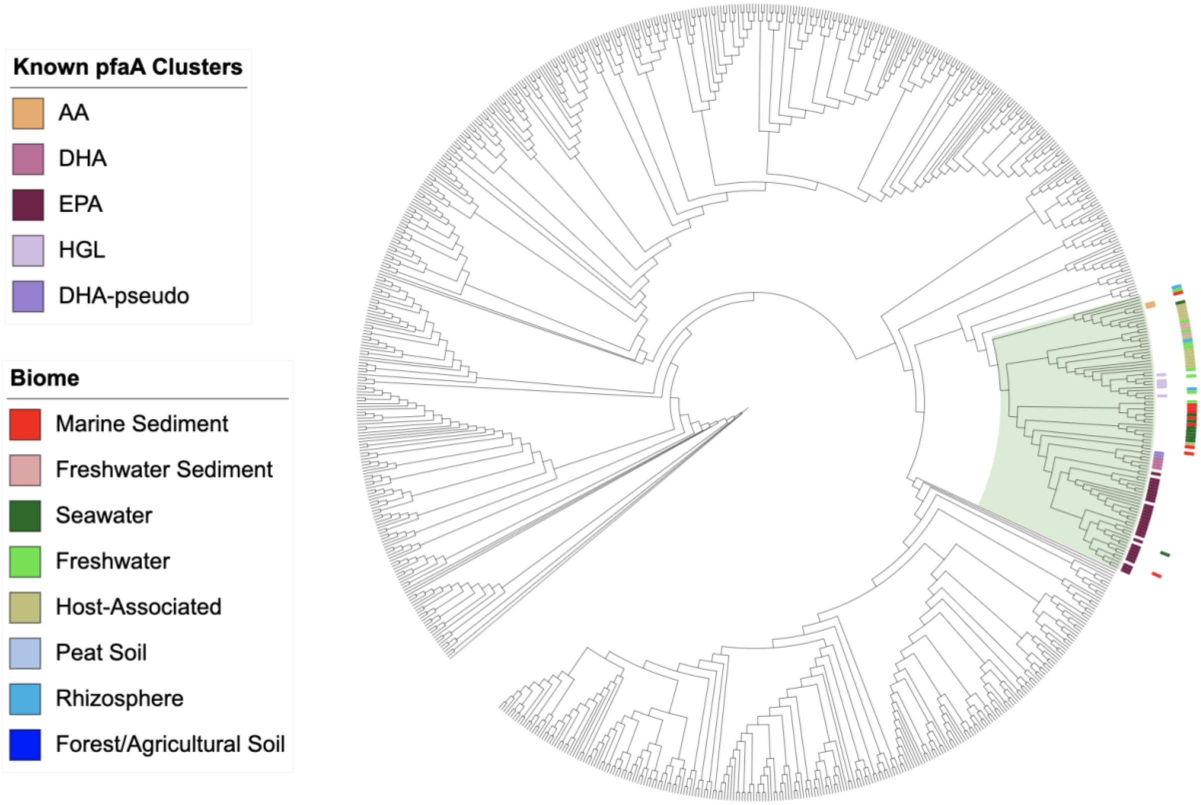


**Figure S10 - pfaA KS domain phylogeny compared to previously characterized pfaA KS clusters.** FastME phylogeny generated from all full-length metagenome-extracted PUFA KS domains that were further classified as belonging to a pfaA module (n=1170) compared against previously characterized pfaA clusters (22). The clade in green includes all pfaA KS domains from these five previously characterized clusters - AA, DHA, EPA, HGL and pseudo-DHA, with 92% of all pfaA KS domains falling outside this clade.

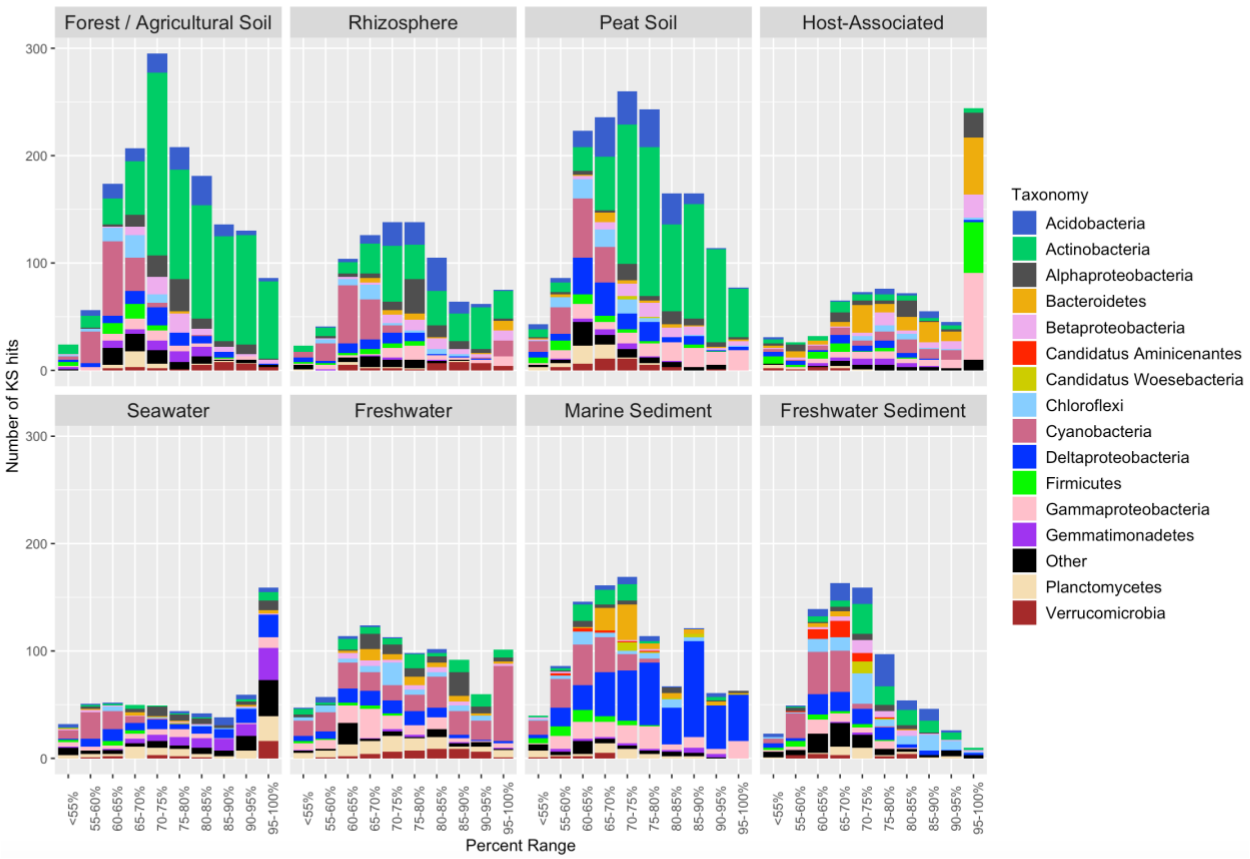


**Figure S11 - Full-length KS domin distributions across biomes.** Stacked bar charts indicate the phylum-level taxonomic composition and abundance (y-axis) of full-length metagenome-extracted KS domains for eight biomes grouped based on percent identity with the closest NCBI BLASTp database match (x-axis).

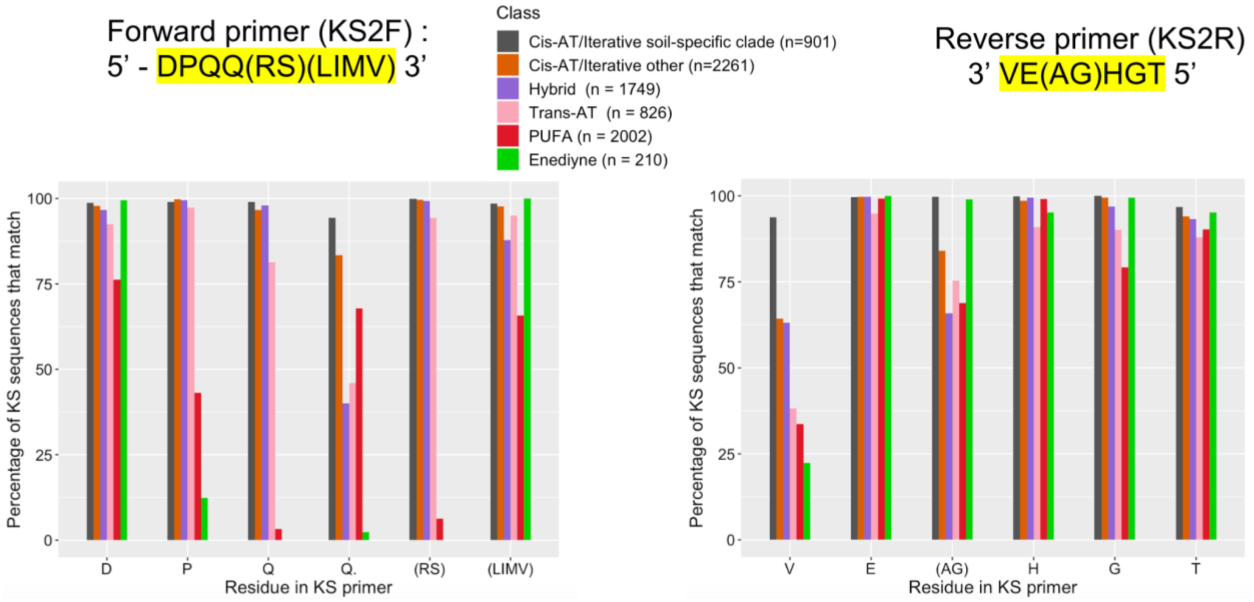


**Figure S12 - Evaluation of the KS2F/R primer set.** Percentage of metagenome extracted KS sequences (y-axis) that match the KS2F/R primers at each amino acid position. KS sequences are grouped by their NaPDoS2 classification. The *cis*-AT/iterative soil-dominant KSs (black) were analyzed separately from all other *cis*-AT/iterative KSs (orange).

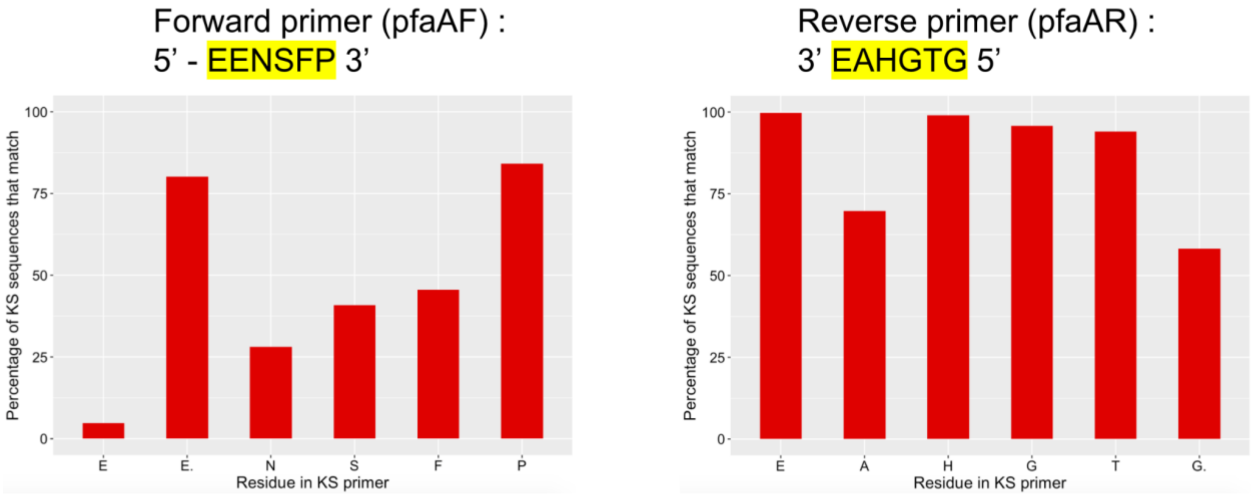


**Figure S13 - Evaluation of the pfaA primer set.** Bar charts were used to visualize the percentages (y-axis) with which the amino acid residues (x-axis) from the PUFA-specific pfaAF/R primer set matched with metagenome-extracted PUFA KS domains (n=1170) identified by NaPDoS2.
